# Ubx5-Cdc48 assists the protease Wss1 at DNA-protein crosslink sites in yeast

**DOI:** 10.1101/2022.05.30.493988

**Authors:** Audrey Noireterre, Ivona Bagdiul, Nataliia Serbyn, Françoise Stutz

**Affiliations:** Department of Molecular and Cellular Biology, University of Geneva, 1211 Geneva 4, Switzerland; Department of Cell Biology, Harvard Medical School, Boston, USA

**Keywords:** Yeast, DNA-protein crosslink, Ubx5, Cdc48/p97, Wss1, Ddi1, UBX, Rpb1

## Abstract

DNA-protein crosslinks (DPCs) pose a serious threat to genome stability. The yeast proteases Wss1, 26S proteasome and Ddi1 are safeguards of genome integrity by acting on a plethora of DNA-bound proteins in different cellular contexts. The AAA ATPase Cdc48/p97 is known to assist Wss1/SPRTN in clearing DNA-bound complexes, however its contribution to DPC proteolysis remains unclear. Here, we show that the Cdc48 adaptor Ubx5 is detrimental in yeast mutants defective in DPC processing. Using an inducible site-specific crosslink, we show that Ubx5 accumulates at persistent DPC lesions in the absence of Wss1, which prevents their efficient removal from the DNA. Abolishing Cdc48 binding or complete loss of Ubx5 suppresses the sensitivity of *wss1Δ* cells to DPC inducing agents by favoring alternate repair pathways. We provide evidence for cooperation of Ubx5-Cdc48 and Wss1 in the genotoxin-induced degradation of RNAPII, a described candidate substrate of Wss1. We propose that Ubx5-Cdc48 assists Wss1 for proteolysis of a subset of DNA-bound proteins. Together, our findings reveal a central role for Ubx5 in DPC clearance and repair.

## INTRODUCTION

DNA-protein crosslinks (DPCs), also known as protein-DNA adducts, are formed by the covalent association of a protein with DNA. If left unresolved, these highly mutagenic and cytotoxic lesions can interfere with essential DNA transactions, cause a severe block to the progression of replication and transcription machineries, and therefore jeopardize the fidelity of genome integrity (Stingele, Bellelli, and Boulton 2017). DPCs are classified into two groups, non-enzymatic and enzymatic, depending on the nature of the crosslinked protein and the triggering mechanism that leads to DPC formation. Non-enzymatic DPCs arise from non-specific crosslinking after exposure to metabolic products such as formaldehyde or acetaldehyde or by the action of exogenous agents (UV-light, IR, etc.), while enzymatic DPCs are the result of abortive enzymatic reactions that require the establishment of a covalent DNA-enzyme reaction intermediate (Stingele and Jentsch 2015; Stingele, Habermann, and Jentsch 2015; Vaz, Popovic, and Ramadan 2017; Zhang, Xiong, and Chen 2020).

DNA Topoisomerase 1 (Top1) is known to be prone to enzymatic DPC formation, as it forms a covalent link with DNA to relax torsional stress that emerges during DNA replication and transcription (Pommier et al. 2016). To do so, Top1 covalently binds one DNA strand to assemble in a transient entity known as a Top1 cleavage complex (Top1cc) (Pommier 2006). Top1-DNA crosslinks can appear when the enzymatic cleavage occurs close to certain DNA lesions (abasic sites, DNA breaks, base mismatch, etc.) or be induced by camptothecin (CPT), a Top1-poison that inserts into the catalytic pocket of Top1 and stabilizes Top1ccs (Staker et al. 2002). Similarly, Flp recombinase introduces a nick at a specific Flp Recognition Target (*FRT*) target site. A Flp mutant carrying the point mutation H305L will drive the formation of a stable covalent crosslink upon *FRT* cleavage. The genetic requirements for cells to repair this DPC-like lesion on DNA are the same as for cells treated with the Top1-poison CPT (Nielsen et al. 2009; Serbyn et al. 2020; Serbyn et al. 2021).

Along with the canonical double-strand breaks (DSBs) repair pathways such as nucleotide excision repair (NER) and homologous recombination (HR) (Ide et al. 2011), numerous other pathways have been implicated in the processing of trapped-Top1ccs (Pommier et al. 2014; Vaz, Popovic, and Ramadan 2017; Ide et al. 2018; Fielden et al. 2018). The tyrosyl-DNA phosphodiesterase Tdp1, which directly hydrolyzes the bond between Top1 and the DNA, was among the first DNA repair enzymes described to have relevance for the elimination of Top1-DNA crosslinks (Yang et al. 1996; Pouliot et al. 1999; Pouliot, Robertson, and Nash 2001). Tdp1 is not able to process intact Top1ccs and prior proteolytic digestion or denaturation is essential to enable hydrolysis by Tdp1 (Yang et al. 1996; Debethune et al. 2002; Pommier et al. 2014). Recently, yeast proteases Wss1 (SPRTN or DVC1 in mammals), Ddi1, and 26S proteasome were shown to participate in the proteolysis of the protein moiety of a DPC (Stingele et al. 2014; Serbyn et al. 2020). It is proposed that remodeling or initial processing of the adducts is required before proteolysis, which may involve the AAA ATPase Cdc48 (known as p97/VCP) (Nie et al. 2012; Stingele et al. 2014; Balakirev et al. 2015; Fielden et al. 2020).

Cdc48/p97 is an abundant essential chaperone involved in a wide variety of cellular processes, including but not limited to cell cycle regulation, membrane fusion, transcriptional control, and ubiquitin-dependent protein degradation (Woodman 2003; Ye 2006; White and Lauring 2007). More recently, several studies investigated its role in DNA replication (Yamada et al. 2000; Mouysset et al. 2008; Deichsel, Mouysset, and Hoppe 2009; Ramadan et al. 2017) and DNA repair (Zhang et al. 2000; Vaz, Halder, and Ramadan 2013; Torrecilla, Oehler, and Ramadan 2017), including DPC repair. In most of the processes Cdc48/p97 processing is initiated by unfolding of ubiquitin conjugated to its targets (Ye 2006; Twomey et al. 2019). The substrate specificity is achieved through interaction with numerous regulatory adaptors (Hanzelmann and Schindelin 2017), of which UBX proteins constitute the largest known group (Schuberth and Buchberger 2008). Supporting the idea of a role for Cdc48/p97 in DNA-bound protein clearance, Cdc48/p97 mediates chromatin extraction and disassembly of some protein complexes (Jentsch and Rumpf 2007; Shcherbik and Haines 2007; Maric et al. 2014; Ramadan et al. 2017; Frattini et al. 2017). It was linked to processes involved in SPRTN/Wss1 proteolysis (Nie et al. 2012; Davis et al. 2012; Stingele et al. 2014; Balakirev et al. 2015; Maskey et al. 2017), and is also required for proteasome-mediated degradation of stalled RNA Polymerase II (RNAPII) following UV-induced DNA damage (Verma et al. 2011; Lafon et al. 2015; He et al. 2017). Particularly, a specialized complex containing mammalian p97 and its cofactor TEX264 is required for clearance of Top1-DNA adducts by SPRTN (Fielden et al. 2020), raising the question of cofactor specificity regarding DPC substrates. The ability of Cdc48/p97 to counteract DPCs appears crucial, however, the mechanistic basis of how it is achieved is still not fully defined (Lin et al. 2008; Mosbech et al. 2012; Nie et al. 2012; Stingele et al. 2014; Balakirev et al. 2015; Fielden et al. 2020).

In this work, we investigate the involvement in DPC repair of Ubx5, a Cdc48 cofactor of the UBX protein family that has previously been implicated in UV-induced degradation of the largest subunit of RNAPII, Rpb1 (Verma et al. 2011). Based on our findings, Ubx5 has a negative effect in *Saccharomyces cerevisiae* (*S. cerevisiae*) mutant strains exhibiting DPC persistence related to the unavailability of the DPC-protease Wss1. By using an *in vivo* inducible DPC system, we observed that Ubx5 accumulation at sites of unrepaired DPC lesions delays their clearance. We further demonstrate that loss of interaction between Ubx5 and Cdc48 is sufficient to suppress the growth phenotype and drug sensitivity of yeast mutant strains accumulating DPCs. Finally, loss of Ubx5 restores the impaired genotoxin-induced degradation of Rpb1 in the absence of the protease Wss1, in a Ddi1-dependent manner. We propose a new mechanism in which Ubx5 mediates Cdc48 recruitment to persistent adducts on chromatin, such as DNA-protein crosslinks or stalled RNA Polymerase II.

## RESULTS

### Multiple Cdc48 cofactors suppress effects of mutations affecting Top1-DNA crosslinks

Aiming to unravel Cdc48 adaptors that define substrate-specificity in DPC repair, we intersected the list of known *CDC48* interactors with the genetic network of the *tdp1-degron + auxin wss1Δ* (*tdp1wss1)* mutant defective in Top1cc repair. A list of physical and genetic *CDC48* interactors was extracted from the yeast Saccharomyces Genome Database [SGD; (Cherry et al. 1997)] (Figure S1A). For these selected genes, we then analyzed the relative number of transposition events in the auxin-treated *tdp1-degron wss1Δ* mutant [Figure 1A; (Serbyn et al. 2020)]. We noted increased transpositions in the *UBX5* gene body (Figure 1B) and confirmed that *ubx5Δ* substantially rescues the growth phenotype of freshly dissected yeast spores (Figure 1C), as well as the CPT sensitivity (Figure 1D) of the *tdp1wss1* double mutant defective in Top1ccs repair (Stingele et al. 2014; Balakirev et al. 2015). It is worth pointing out that Ufd1 and Npl4, both known to be core adaptors of the Cdc48-Ufd1-Npl4 complex recruited to DSBs (Meerang et al. 2011) and DNA-bound proteins (Balakirev et al. 2015), are suppressors of *tdp1wss1* as well (Figure 1A). Moreover, Ubx5 protein levels are decreased in the *tdp1wss1* mutant, emphasizing the negative effect of Ubx5 in that cellular context (Figures 1E and 1F; *tdp1wss1* + auxin). Loss of Wss1 alone already causes reduction of steady-state Ubx5 levels (Figures 1E and 1F; *tdp1wss1* no auxin), suggesting that the *ubx5Δ* suppression may represent the adaptive response to Wss1 deficiency. From these data, we presumed that Ubx5 may have an inhibitory effect in the absence of the repair enzymes Wss1 and Tdp1.

**Figure 1.**
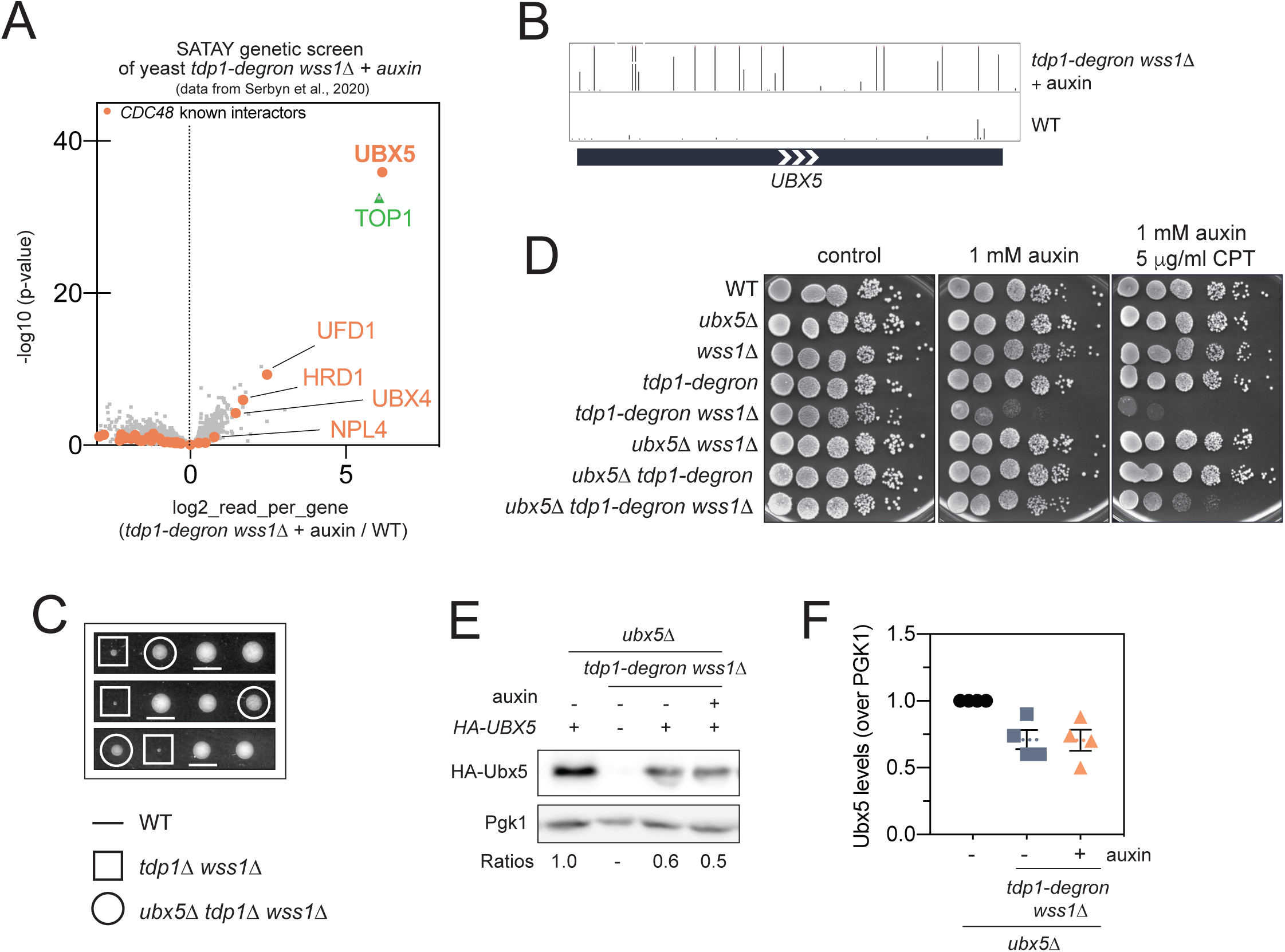
Cdc48 cofactor Ubx5 is a new suppressor of *tdp11Δ wss11Δ*. (A) *CDC48* known interactors (orange) identified as suppressors or sensitizers in the *tdp1-degron wss1*1Δ SATAY transposon screen described in (Serbyn et al. 2020). The volcano plot compares the sequencing reads in *tdp1-degron wss1*1Δ + auxin and a pool of six unrelated SATAY libraries. Fold-change of reads per gene (log2, x-axis) and corresponding p-values (-log10, y-axis) are plotted. The gene names of the strongest suppressors are labelled in orange. Top1 (positive control) appears in green. (B) Snapshot depicting transposon coverage of *UBX5* gene body in the *tdp1-degron wss1*1Δ + auxin and one of the WT libraries. The height of the bars represents the number of reads for each transposon. (C) Loss of *UBX5* suppresses growth defects of *tdp11Δ wss11Δ*. Tetrads were analyzed after dissection of the diploid [*TDP1/tdp1*1Δ; *WSS1/wss1*1Δ; *UBX5/ubx5*1Δ]. (D) Transposon screen validation of *UBX5. ubx5*1Δ restores the growth of *tdp1wss1*. Cells were grown in YEPD and spotted on a medium supplemented with 1 mM auxin and 5 μg/mL Camptothecin (CPT). Plates were incubated for 2 days at 30°C. (E-F) Ubx5 levels are decreased in *tdp1wss1.* (E) HA-Ubx5 protein levels at steady state are compared by immunoblotting in different mutants. Mutant strains carrying additional *ubx51Δ* mutations were complemented with a plasmid expressing Ubx5 from the endogenous *pUBX5* promoter. 1mM auxin was added where indicated for 6 h to degrade *tdp1-degron* (Tdp1-AID*-9MYC), when present. Pgk1 was used as a loading control. For quantification (F), the anti-HA signal was compared to Pgk1 and normalized to WT. Quantification of 4 biological replicates are presented as means ± SEM.

### Interaction between Cdc48 and Ubx5 interferes with the resistance to DPC-inducing agents in the absence of the Wss1 protease

Wss1 protease is required for cellular resistance against formaldehyde (FA) and hydroxyurea (HU) [Figure 2A, (O’Neill et al. 2004; Maddi et al. 2020; Serbyn et al. 2020)]. While it is well-documented that HU induces replication stress, several studies reported that HU creates free nitroxide radicals (Yarbro 1992; King 2003), which are prone to create DPCs (Dizdaroglu et al. 1989; Nakano et al. 2003), raising the possibility that DPCs may increase in response to HU. Since Ubx5 levels are downregulated in the absence of Wss1 (Figures 1E and 1F), we hypothesized that Ubx5 loss could restore tolerance to these DPC-inducing agents. Indeed, *ubx5Δ* reverts FA and HU hypersensitivity of the *wss1Δ* mutant to WT-like levels (Figure 2A). In contrast, *ubx5Δ* did not suppress hypersensitivity to Top1ccs in the absence of Tdp1 (Figure S2A). From these data, we inferred that Ubx5 is genetically linked to Wss1 but not to the Top1-specific repair factor Tdp1. Similarly to Wss1, Ubx5 might target a broad range of DPCs for repair.

**Figure 2.**
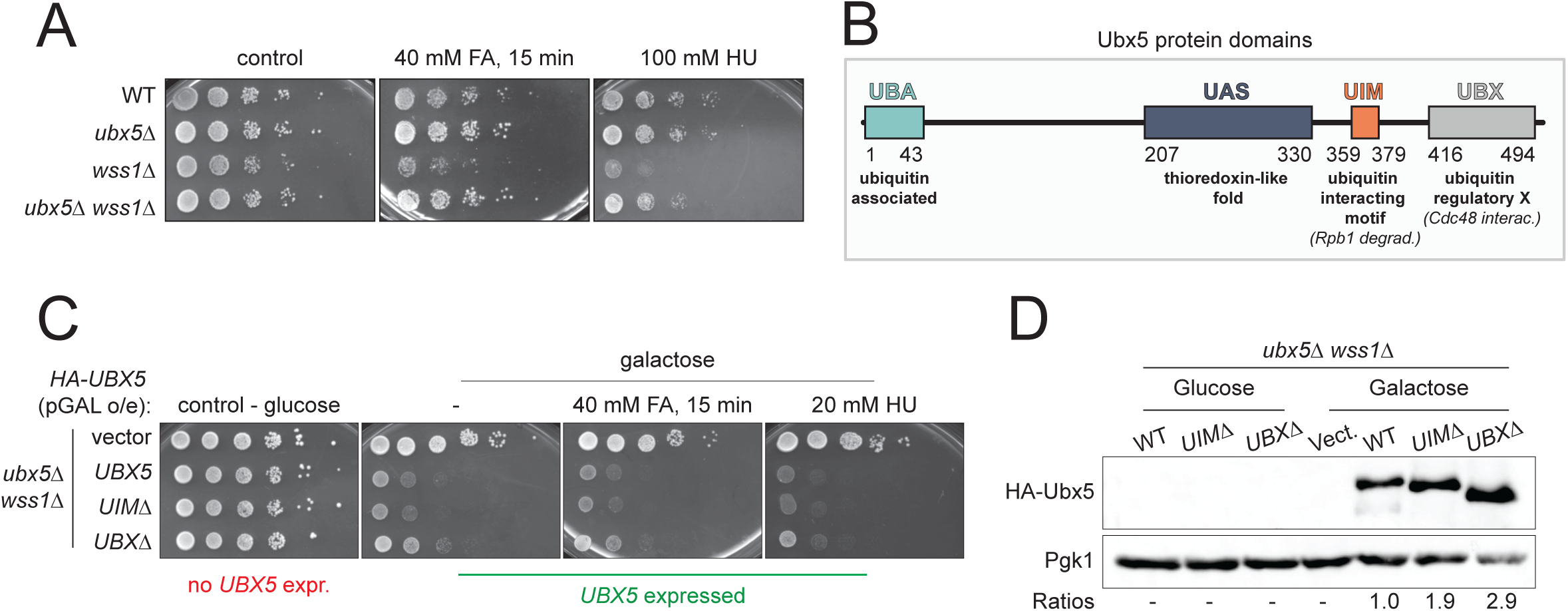
Loss of interaction between Cdc48 and Ubx5 restores growth of cells accumulating Top1ccs. (A) Ubx5 loss rescues *wss1*1Δ sensitivity to formaldehyde (FA) and hydroxyurea (HU). Cells were grown in YEPD and either treated with 40 mM FA for 15 min or spotted directly on 100 mM HU plates and incubated for 2 days at 30°C. (B) Schematic representation of Ubx5 domains. Yeast Ubx5 consists of four known domains: an amino (N)-terminal UBA, central UAS, UIM, and a carboxyl (C)-terminal UBX. UBX domain is a Cdc48 binding module. UIM is implicated in Rpb1 degradation after UV treatment. (C) *UBX*1Δ mutant of Ubx5 is sufficient to suppress FA and HU sensitivities of *wss1*1Δ. *ubx51Δwss11Δ* mutants were complemented with plasmids encoding HA-Ubx5-WT or HA-Ubx5 mutants overexpressed under the control of the *pGAL* promoter; cells were grown in SC-trp and either treated with 40 mM FA for 15 min or spotted directly on 20 mM HU. Plates were prepared either with glucose (no HA-Ubx5 expression) or galactose (Ubx5 expression) as the source of sugar. Plates were incubated for 3 days at 30°C. o/e, overexpression. (D) Protein levels of overexpressed HA-Ubx5 mutants shown in (C) were analyzed by immunoblotting. For quantification, the anti-HA signal was compared to Pgk1 and then normalized to HA-Ubx5-WT expressed in galactose.

To gain further insights into Ubx5 function in repair, we addressed the role of Ubx5 domains. Yeast Ubx5 consists of four known domains: an amino (N)-terminal UBA, central UAS, UIM, and a carboxyl (C)-terminal UBX (Figure 2B). To test the role of some Ubx5 domains, we complemented *ubx5Δ wss1Δ* with HA-tagged Ubx5 constructs overexpressed from a galactose-inducible promoter (Figure 2C). Complementation by *ubx5^uimΔ^* was detrimental for cell growth and compromised resistance to several drugs, similar to overexpression of the full-length protein, emphasizing the fact that the UIM domain is not responsible for the negative effect of Ubx5 in the absence of Wss1. This observation contrasts the previously documented role of UIM in RNAPII turnover after UV treatment (den Besten et al. 2012), which implicated Ubx5 in DNA damage response. We next assayed whether the UBX domain that mediates the interaction with Cdc48 (Schuberth and Buchberger 2008) is critical for DPC repair. Overexpression of the *ubx5^ubxΔ^* mutant showed partial rescue of *ubx5Δ wss1Δ* phenotype, even in the presence of FA and HU (Figure 2C), suggesting the importance of Ubx5-Cdc48 interaction. Of note, this is not due to a reduced protein level of the *ubx5^ubxΔ^* variant (Figure 2D). Ubx4 is another UBX-containing protein (Schuberth and Buchberger 2008) identified as a mild suppressor of *tdp1wss1* (Figure 1A) that presents a highly transposed UBX domain in the screen (Figure S2B). The *ubx4^ubxΔ^* mutant partially rescues HU sensitivity of *wss1Δ* cells, as well as the growth defect of *tdp1wss1* (Figures S2B and S2C) suggesting that Ubx4 could function as an alternative Cdc48 adaptor needed for DPC repair. Taken together, these findings support the view that Ubx5 and Ubx4, both cofactors of Cdc48, are detrimental upon DPC accumulation through their interaction with the segregase Cdc48 via their UBX domain.

### Loss of the Cdc48 cofactor Ubx5 improves repair efficiency of the crosslinked Flp

Ubx5 mutants restore a WT-like phenotype in the absence of the repair enzymes Wss1 and Tdp1. Accumulation of Top1 crosslinked to DNA in *tdp1Δ wss1Δ* was previously demonstrated (Stingele et al. 2014). We reasoned that loss of *UBX5* would lead to decreased number of crosslinked proteins detected on DNA. To test this hypothesis, we used the previously described Flp-nick system [Figure 3A; (Nielsen et al. 2009)], which creates a Top1-like crosslink at a single *FRT* locus artificially introduced into chromosome VI of the yeast (*S. cerevisiae)* genome. This system uses a galactose-inducible mutant Flp recombinase *flp-H305L,* which triggers a “Flp-cc” at the *FRT* site, thus allowing to precisely follow the molecular events linked to the appearance, detection and repair of the Flp-cc. Similar to the genetic interactions observed in the *W303* strain background (Figure 1D), *ubx5Δ* also suppressed cell growth defects resulting from *flp-H305L* expression on 2% galactose in the Flp-nick system (Figure 3B). Of note, as the Flp-nick system mimics a Top1cc, endogenous Top1 was removed in the following experiments.

**Figure 3.**
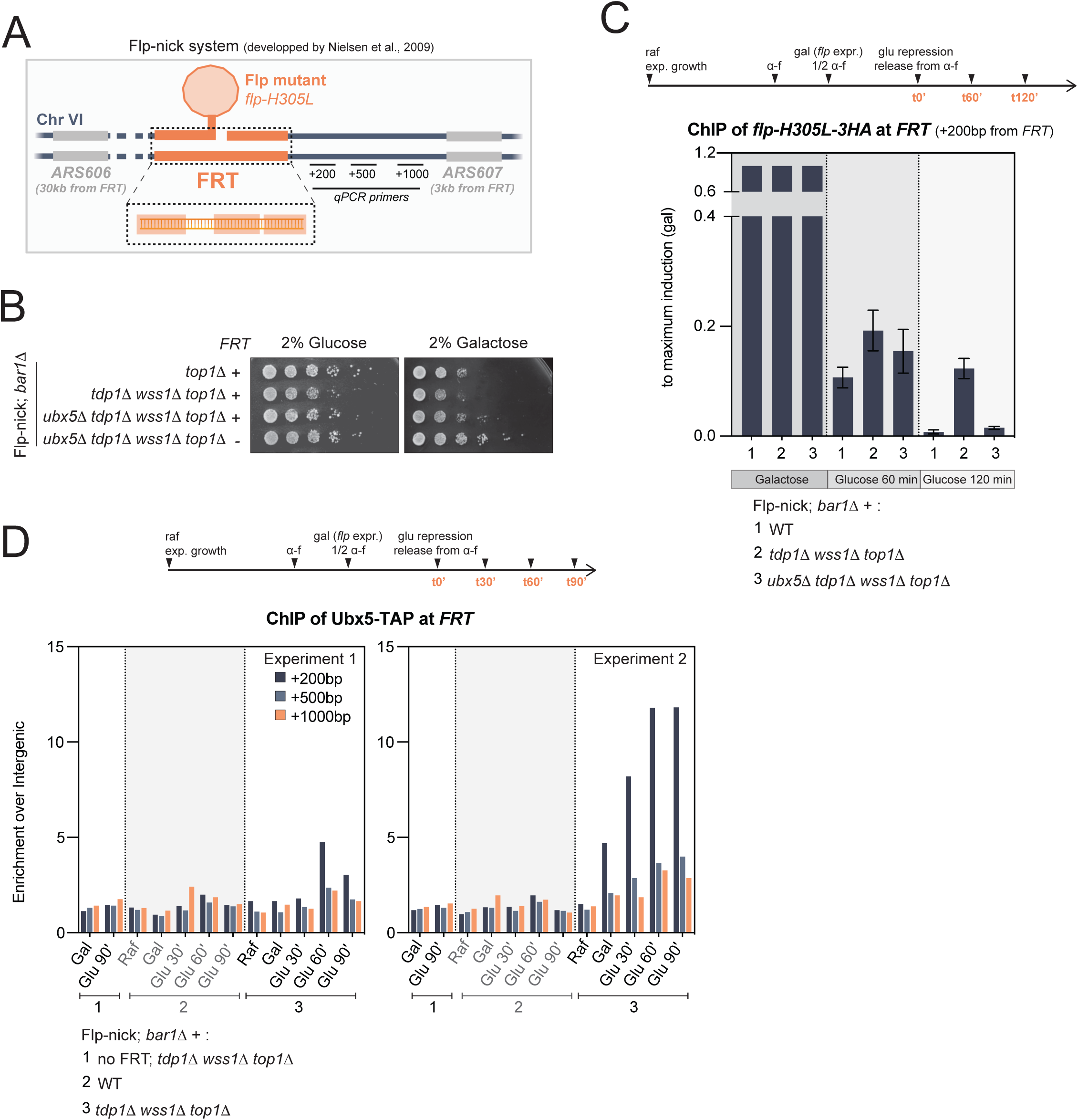
Ubx5 has a suppressing effect on crosslinked Flp. (A) Schematic representation of the “Flp-nick” system initially described in (Nielsen et al. 2009). The mutant *flp-H305L* recombinase is expressed from a galactose-inducible promoter and targeted to the *FRT* site introduced in the yeast genome next to *ARS607*, on chromosome VI. *FRT* consists of three DNA elements (orange) recognized by the Flp recombinase. The location of qPCR primers used for all subsequent ChIP-qPCR analyses is indicated in black, at +200bp, +500bp, and +1000bp from the *FRT*. (B) The *ubx5*1Δ mutation rescues growth defects caused by Flp-nick galactose-induction in *tdp1*1Δ *wss1*1Δ *top1*1Δ mutant. Indicated strains were grown in YEP-2% raffinose prior to plating on YEP-2% glucose or YEP-2% galactose plates. Plates were incubated for 3 days at 30°C. (C) *flp-H305L-3HA* dynamics at the *FRT* site in different mutants. Cells were grown in 2% raffinose (raf), synchronized in G1 with α-factor (α-f) during induction of *flp-H305L* expression with 3% galactose (gal). Induction was stopped by addition of glucose (glu) and cells were released into the cell cycle. Samples were collected at the indicated time points. Levels of *flp-H305L-3HA* were assessed by ChIP and qPCR without FA crosslinking. The positions of qPCR primers +200bp, +500bp, and +1000bp downstream of *FRT* are indicated in (A). The graph only shows the results with +200 qPCR primers. See Figure S3B for other primers. No FRT binding site and no *flp-H305L*-*3HA* expression (raf) were used as negative controls. The percentage of input for each genotype was normalized to maximum galactose induction. Data are presented as the means ± SEM of two independent replicates. See Figure S3A for cell cycle analyses by FACS. See Figure S3B for *flp-H305L-3HA* levels after maximum galactose-induction. (D) Ubx5 occupancy at the *FRT* site. Levels of Ubx5-TAP were monitored by ChIP and qPCR following FA crosslinking in cells grown as described in (C). Quantifications from two independent experiments are shown. No *FRT* strain and raffinose samples were used as negative controls.

To monitor changes in *flp-H305L* retention at the *FRT* site, cells were first synchronized in the G1-phase of the cell cycle by the addition of α-factor during the 2 h galactose induction of the *flp-H305L* mutant (Figure 3C, upper panel). *flp-H305L* induction (see Figure S3B to compare galactose-induction levels) was then stopped with glucose and cells were released from G1 arrest and allowed to enter the cell cycle (Figure 3C, glu 60 min and 120 min; Figure S3A). Consistently, and as recently reported (Serbyn et al. 2020; Serbyn et al. 2021), *flp-H305L* was more persistent at the *FRT* site at 120 min in *tdp1Δ wss1Δ* compared to a WT strain, which showed rapid removal of the Flp-cc from the *FRT* site (Figure 3C, compare strains 1 and 2). Since *UBX5* deletion suppresses the *tdp1Δ wss1Δ* phenotype, we hypothesized that Ubx5 removal may lead to faster clearance of the protein nick. Indeed, Flp-cc repair dynamics in *ubx5Δ tdp1Δ wss1Δ* were very similar to WT (Figure 3C, strain 3). These results suggest that the rescue observed in *ubx5Δ* is probably due to a reduced amount of DPCs on the DNA.

To define whether the possible negative effect of Ubx5 could be due to its presence at the damage site, we tested Ubx5 recruitment at the *FRT* site using the Flp-nick system. We observed that while Ubx5 was not recruited at the Flp-cc site in a WT-like strain, there was a great enrichment of Ubx5 upon removal of the Wss1 and Tdp1 repair enzymes that was not detected in the absence of *FRT* (Figure 3D). A straightforward interpretation of these findings is that Ubx5 prevents the elimination of the Flp-cc (and probably Top1cc, and other DPCs) from the DNA by hyper-accumulating at the damage location when Wss1 and Tdp1 are unavailable.

### Other repair pathways are active in *ubx5Δ tdp1Δ wss1Δ*

We postulated that *UBX5* deletion could lead the cells towards an alternative repair pathway. The aspartic protease Ddi1 has recently been characterized as a novel DPC repair player acting in parallel to Wss1 (Svoboda et al. 2019; Serbyn et al. 2020). We thus speculated that Ddi1 could play a role in improving the growth of *ubx5Δ tdp1Δ wss1Δ*. By performing genetic analyses, we observed that Ddi1 was indeed essential for *ubx5Δ tdp1Δ wss1Δ* survival, as spores additionally lacking Ddi1 were unable to proliferate (Figure 4A). Likewise, Ddi1 was crucial for the viability of the *ubx5Δ wss1Δ* mutant under HU stress (Figure S4A).

**Figure 4.**
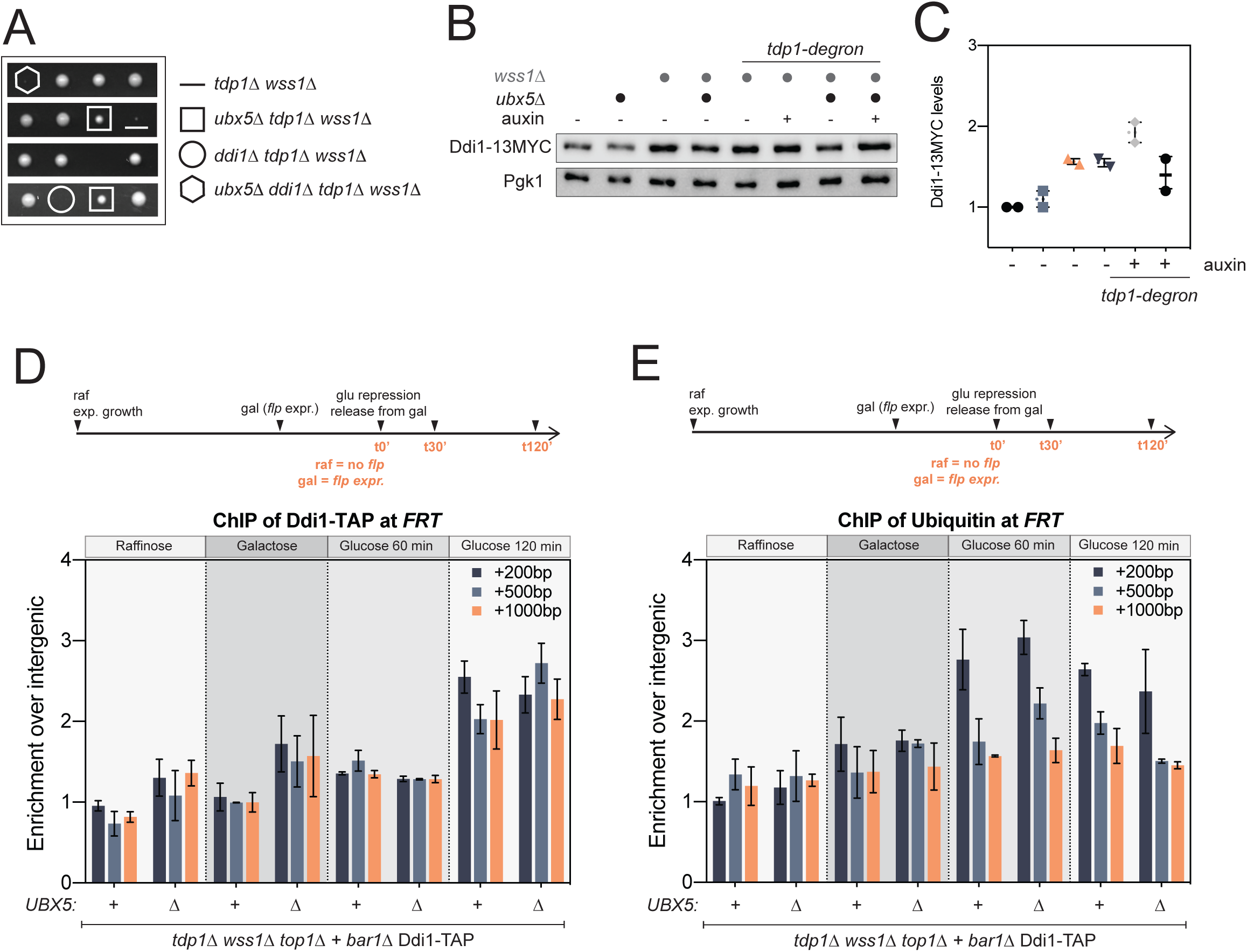
The suppressing effect of *ubx51Δ* partially depends on the protease Ddi1. (A) The suppressing effect of *ubx5*1Δ in *tdp1*1Δ *wss1*1Δ depends on the ubiquitin-dependent protease Ddi1. Tetrads were analyzed after dissection of the diploid [*TDP1/tdp1*1Δ; *WSS1/wss1*1Δ; *UBX5/ubx5*1Δ; *DDI1/ddi1*1Δ]. See also Figure S4A for HU sensitivities. (B) Endogenous Ddi1-13MYC levels are still elevated upon additional *ubx5*1Δ mutation. 1mM auxin was added where indicated for 6 h to degrade *tdp1-degron* (Tdp1-AID*- 6HA), when present. Levels in total cell extracts were compared by immunoblotting. (C) Quantification of Ddi1-13MYC levels shown in (B). The anti-MYC signal was compared to Pgk1 and normalize to WT. Quantification of 2 biological replicates are presented as means ± SEM. (D) Ddi1 occupancy at the *FRT* site. Levels of Ddi1-TAP were examined by ChIP-qPCR following formaldehyde crosslinking. Cells were grown as explained in legend of Figure 3C, without α-factor synchronization, and collected at the indicated time points. The enrichments relative to intergenic were normalized to raffinose (+200) and plotted ± SEM of two independent replicates. (E) Ubiquitination of the *FRT* locus. Asynchronous cultures were grown in 2% raffinose (raf) or additionally supplemented with 3% galactose (gal) to induce *flp-H305L* expression. Ubiquitin antibody was used for ChIP-qPCR analysis following FA crosslinking. Data are shown as the mean ± SEM of two independent replicates. Enrichments at the *FRT* locus were normalized to raffinose (+200).

We further tested the requirements of other known Top1ccs and DPC repair factors (Liu, Pouliot, and Nash 2002; Deng et al. 2005; Pommier 2006; Alvaro, Lisby, and Rothstein 2007; Stingele et al. 2014; Sun et al. 2020) (Figures S4B to S4K). Among the pathways tested, deletion of *RAD52* (HR, Figure S4B), *RAD9* (checkpoint activation, Figure S4I), *SGS1* (RecQ helicase, Figure S4G), *SRS2* (anti-recombinase, Figure S4H), *RAD27 and MRE11* (structural nucleases, Figures S4C and S4D), strongly affected growth of *ubx5Δ tdp1Δ wss1Δ* spores, emphasizing the essentiality of these enzymes and their respective repair pathways to counteract DPC appearance in yeast mutants lacking Wss1 and Tdp1. Surprisingly, abrogation of the non-homologous end-joining (NHEJ) pathway (*yku70Δ*, Figure S4E), the translesion synthesis pathway (*rev3Δ*, Figure S4J) as well as deletion of the NER pathway (*rad4Δ*, Figure S4F) had no effect on *ubx5Δ tdp1Δ wss1Δ* spore survival. Taken together, these genetic analyses (Figure S4K) show that checkpoint pathways, recombination, multiple nucleases and Ddi1 protease facilitate the growth of *ubx5Δ tdp1Δ wss1Δ*.

### Ddi1 provides resistance towards DPCs to cells lacking Ubx5 and Wss1

Because the Ddi1 and Wss1 proteases work in parallel pathways, Ddi1 becomes more important when Wss1 is absent. Thus, Ddi1 expression is elevated in cells lacking Wss1 (Figures 4B and 4C). Ddi1 protein levels remained overexpressed following deletion of *UBX5* on top of *wss1Δ* mutation, indicating that it still plays a relevant role to help both the *ubx5Δ wss1Δ* and the *ubx5Δ tdp1Δ wss1Δ* mutants in dealing with different sources of stress (Figures 1D and 2A). Accordingly, ChIP-qPCR analyses at the *FRT* locus revealed that Ddi1 is recruited in *tdp1Δ wss1Δ top1Δ* (Figure 4D), as previously observed (Serbyn et al. 2021). Even though the *ubx5Δ tdp1Δ wss1Δ top1Δ* mutant demonstrated more efficient Flp-cc elimination from the *FRT* (Figure 3C), Ddi1 was recruited to the same extent. Since Ddi1 was recently shown to be a ubiquitin-dependent protease (Yip, Bodnar, and Rapoport 2020), we also examined ubiquitination of the DPC-site in *ubx5Δ.* In conjunction with Ddi1 recruitment at the locus, we observed similar accumulation of ubiquitin at the *FRT* site in both *tdp1Δ wss1Δ top1Δ* and *ubx5Δ tdp1Δ wss1Δ top1Δ* (Figure 4E). This enrichment of Ddi1 suggests that it is a key contributor in *ubx5Δ-*mediated suppression of *tdp1Δ wss1Δ*, probably by mediating proteolytic digestion of the DPC.

### Ubx5 prevents efficient turnover of stalled RNAPII in the absence of Wss1

Several studies have revealed that the largest subunit of RNAPII, Rpb1, is prone to stalling and degradation after exposure to genotoxic agents (Beaudenon et al. 1999; Malik et al. 2008; Verma et al. 2011; Wilson et al. 2013). The UV-induced degradation of Rpb1 depends on Cdc48, as well as its adaptor proteins Ubx4 and Ubx5 (Verma et al. 2011). Moreover, stalled Rpb1 degradation after UV or HU exposure also engages the DPC-proteases Ddi1 and Wss1 (Serbyn et al. 2020). It is still unclear how Ddi1 and Wss1 may assist Ubx4-Ubx5-Cdc48 in the extraction of stalled Rpb1, but it is likely that stalled Rpb1 is a substrate of these two proteases.

In light of the suppressing effect of *ubx5Δ* on *wss1Δ* cells and the requirement of Ddi1 in this context, we aimed to assess how Rpb1 stability was altered in these conditions. We examined Rpb1 protein levels in total yeast cell extracts subjected to HU treatment. As formerly reported, Rpb1 was rapidly degraded in response to HU exposure in WT cells, while loss of Ubx5 or Wss1 hindered this process (Figures 5A and 5B). Surprisingly, the double *ubx5Δ wss1Δ* mutant was proficient for Rpb1 turnover, although both proteins alone are strictly required for degradation. Additionally, Rpb1 turnover in *ubx5Δ wss1Δ* was dependent on Ddi1. Indeed, upon HU treatment, Rpb1 degradation was compromised in the absence of Ddi1 (Figures 5A and 5B). These observations reveal that the function of Wss1 in Rpb1 degradation can be bypassed if *UBX5* is deleted, strongly favoring the Ddi1-dependent pathway.

**Figure 5.**
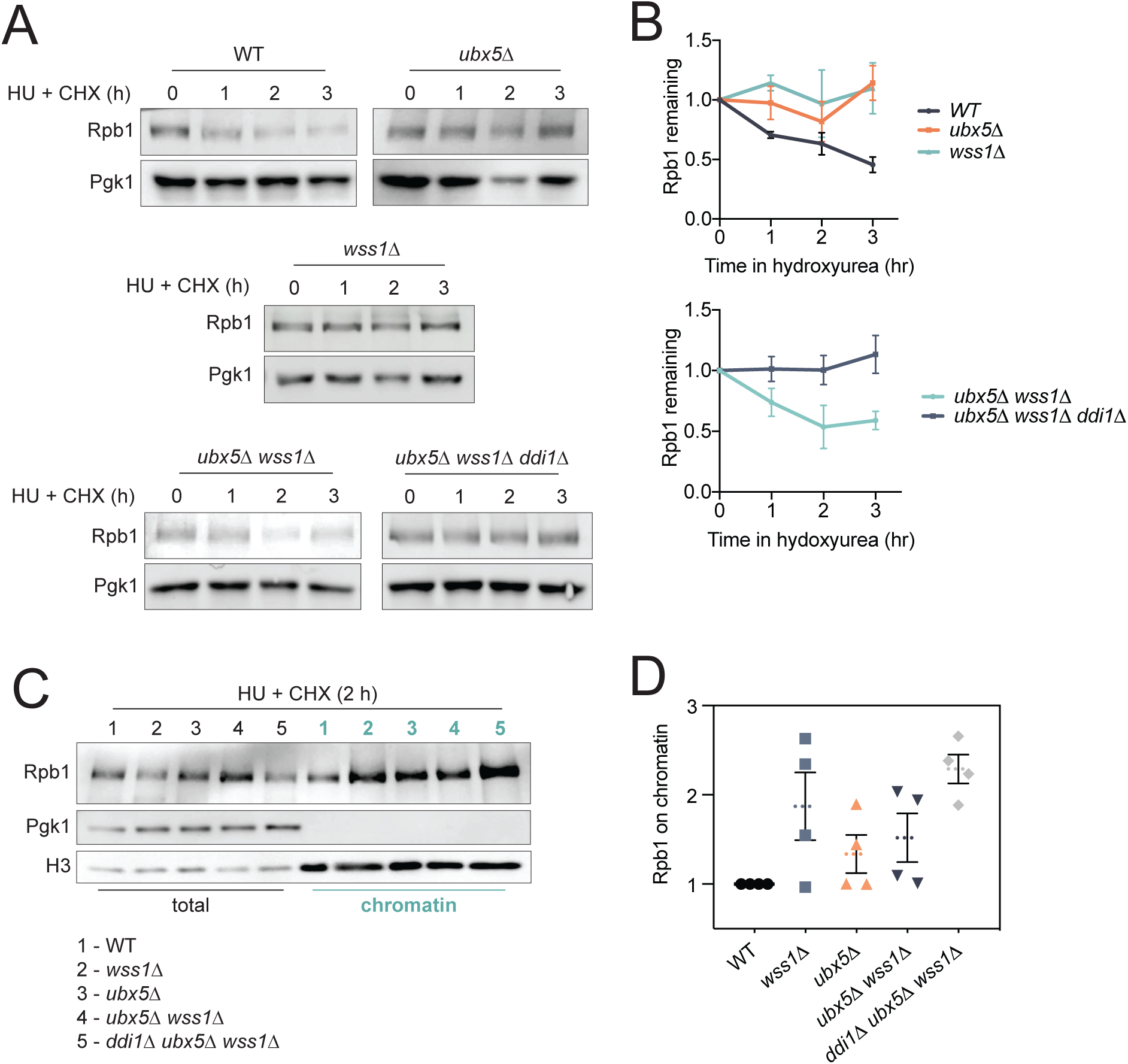
Ubx5 loss promotes Rpb1 degradation in the absence of the DPC protease Wss1. (A) Ubx5 loss restores HU-induced Rpb1 degradation in *wss1*1Δ and it requires Ddi1. Rpb1 levels were defined in cells growing in the presence of 200 mM HU and 100 μg/mL CHX at the indicated time points. Rpb1 and Pgk1 levels in total cell extracts were probed by immunoblotting and quantified using fluorescent secondary antibodies. See (B) for quantifications. Images show immunoblotting using chemiluminescence. (B) Rpb1 turnover representative of pictures shown in (A). Relative Rpb1 to Pgk1 levels were set to 1 in the respective non-treated samples. Graphs show values of three independent replicates (C-D) Rpb1 chromatin enrichment following HU treatment. (C) Chromatin fractions were isolated from cells treated for 2 h with 200 mM HU and 100 μg/mL CHX. Pgk1 and H3 were used as controls to monitor the fractionation. (D) Quantifications of Rpb1 and H3 levels in chromatin fractions. Relative Rpb1 to H3 levels were set to 1 in the respective WT samples. Quantifications of 4 biological replicates are presented as means ± SEM.

With respect to prior work that described Rpb1 accumulation on chromatin in *wss1Δ* (Serbyn et al. 2020), we also examined DNA-bound Rpb1 in the mutants generated in this study. Consistently, Rpb1 accumulation can easily be observed in *wss1Δ* (Figure 5C, lane 2; Figure 5D), and this stabilization is counteracted in *ubx5Δ wss1Δ* (Figure 5C, lane 4). Finally, Rpb1 was highly abundant on chromatin in the triple mutant *ddi1Δ ubx5Δ wss1Δ* (Figure 5C, lane 5; Figure 5D), emphasizing the importance of Ddi1 in the suppressing effect of *ubx5Δ.* Taken together, these data suggest that Ubx5 contribution to the clearance of stalled DNA-bound Rpb1 is relevant only in the presence of Wss1. This argues that Wss1 and Ubx5 may cooperate for genotoxin-induced Rpb1 turnover. When either Wss1 or Ubx5 are lost, the pathway for degradation of stalled RNAPII is compromised and strongly relies on Ddi1.

## DISCUSSION

With the progression of studies on DNA-protein crosslinks, several repair routes have been discovered but the contribution of each pathway still needs to be elucidated. This study aimed to gain a deeper understanding of the involvement of the AAA ATPase Cdc48 in the cellular response to DPCs. Here, we show that the UBX protein Ubx5 is the major Cdc48 adaptor capable of suppressing the phenotype associated with the defective Wss1-dependent route of proteolytic DPC processing. Loss of Ubx5-Cdc48 interaction was sufficient to abrogate phenotypes linked to loss of Wss1 or deletion of the critical Top1cc repair genes *WSS1* and *TDP1*. Our data indicate that Ubx5 is targeted to persistent DPC sites and prevents them from cleavage if Wss1 enzymatic activity is unavailable; they are also consistent with the view that Wss1 cannot efficiently perform its proteolytic activity without prior action of Ubx5-Cdc48, revealing an unexpected role of Ubx5 in coordinating the cooperation between Cdc48 and Wss1 in the removal of DNA-bound proteins. We exploit these observations to propose a model for Ubx5-Cdc48 and Wss1 co-action in DPC elimination (Figure 6), in which Ubx5 targets Cdc48-Ufd1-Npl4 to ubiquitinated DPCs for initial processing of the protein adduct, followed by Wss1 recruitment to complete proteolysis.

**Figure 6.**
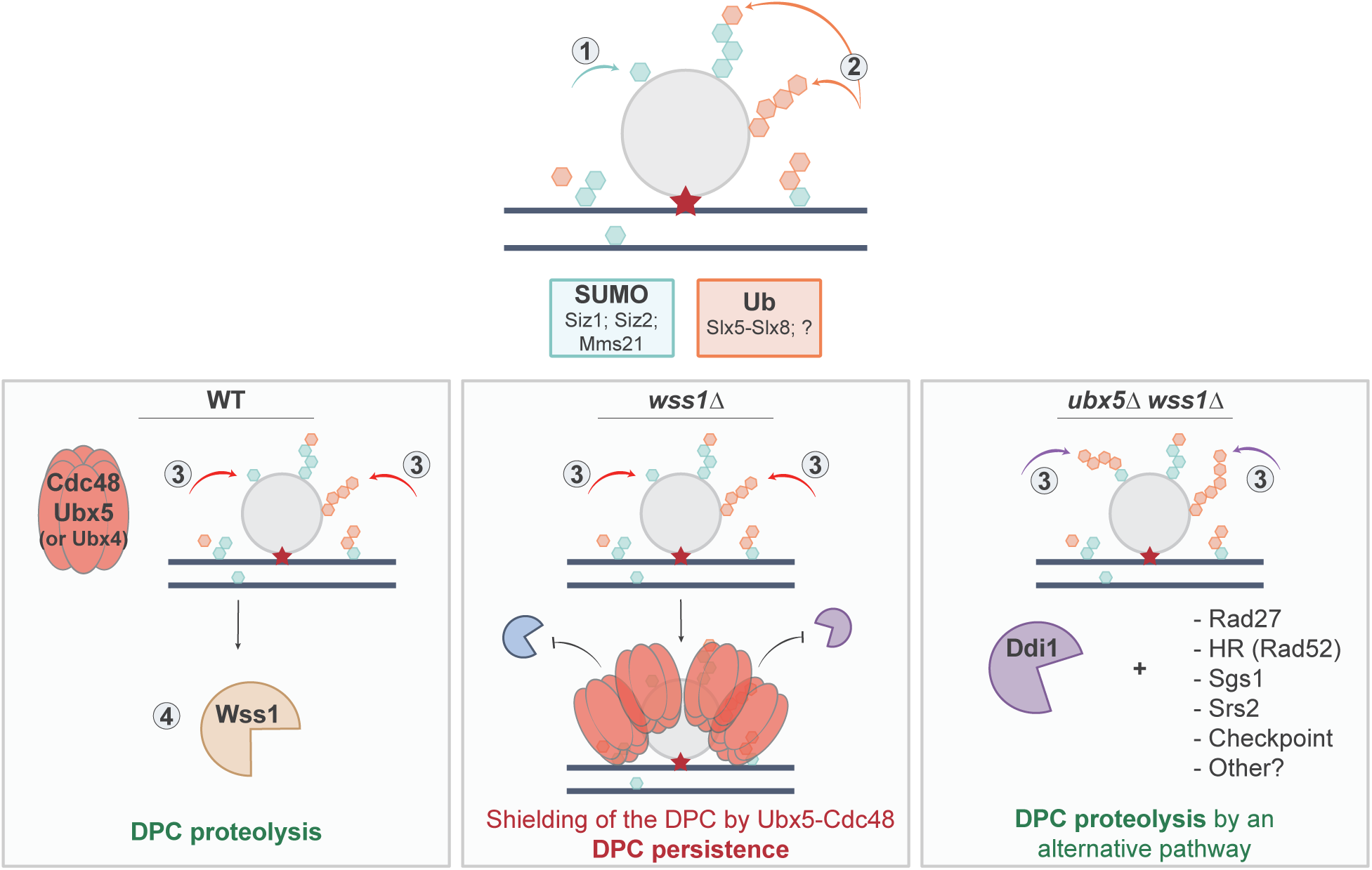
Putative model for Ubx5-Cdc48 in DPC repair. Repair of a DPC (Top1cc, Flp-cc, stalled Rpb1, etc.) in budding yeast *S. cerevisiae*. Persistence of the protein adduct on chromatin results in sumoylation (SUMO; by Siz1, Siz2 or Mms21 E3 ligases) and ubiquitylation (Ub; by Slx5-Slx8 and potentially other E3 ligases), allowing Ubx5-Cdc48-Ufd1-Npl4 loading and initial remodeling of the protein adduct. Proper protein extraction is ensured by the additional recruitment of the protease Wss1. If Wss1 is unavailable (*wss1Δ*), Ubx5-Cdc48 is still targeted to the DPC site, but the proteolysis is not efficient enough to open access for Tdp1 or other endonucleases, and Ubx5-Cdc48 shields the damage site. Accumulation of Ubx5-Cdc48 prevents access to other salvage pathways. Loss of Ubx5 (*ubx5Δ wss1Δ*), or abrogation of its interaction with Cdc48, allows access to the DPC lesion site for alternative repair pathways, which may involve the DPC proteases Ddi1 and 26S proteasome, as well other canonical repair pathways.

### Ubx5-Cdc48 assists Wss1 at DNA-bound proteins

An unexpected finding from our work is that cells deprived of Ubx5-Wss1 are resistant to DPC induction (Figure 2A) and present a normal Rpb1 turnover following exposure to HU (Figures 5A and 5B), although both Ubx5 and Wss1 activities are important to mediate extraction of Rpb1 from chromatin [Figures 5C and 5D; (Verma et al. 2011; Serbyn et al. 2020)]. Interestingly, SPRTN/Wss1 has already been directly connected to p97/Cdc48 in several reports (Davis et al. 2012; Mosbech et al. 2012; Stingele et al. 2014; Balakirev et al. 2015; Fielden et al. 2020; Kroning et al. 2022). Hence, we suggest that the protease Wss1 is assisted by the unfolding activity of Ubx5-Cdc48 in the removal of chromatin-bound proteins. It remains to be elucidated which comes first, Wss1 or Cdc48? On one hand, Cdc48 might extract by-products generated by Wss1 enzymatic activity. On the other hand, the activity of Cdc48 may be required to prepare the DPC and facilitate its proteolysis by Wss1. We envision that Ubx5-Cdc48 is targeted first, as we were able to detect Ubx5 at a Top1cc-like locus in the absence of Wss1 (Figure 3D). Similarly, experiments in human cells showed that p97/Cdc48 enabled proteolysis of Top1cc by SPRTN/Wss1, probably by unfolding the adduct to make it more accessible for SPRTN cleavage (Fielden et al. 2020). We believe that Ubx5-Cdc48 engaged on a permanently crosslinked protein is unable to complete substrate degradation without Wss1. Consequently, in the absence of Wss1, Ubx5-Cdc48 targeting and shielding of the DNA-bound protein will prevent easy access to parallel repair factors, such as Ddi1 (Figure 4). In this view, if Ubx5-Cdc48 is unavailable, the function of Wss1 would also be compromised as the initial processing and unfolding of the adduct cannot take place (Figure 6). Our hypothetical model is also supported by a recent study which demonstrated that the action of p97/Cdc48 was required for SPRTN/Wss1 proteolytic cleavage to occur on tightly bound proteins (Kroning et al. 2022).

### Is Ubx5-Cdc48 strictly required for Wss1 enzymatic activity on DPCs?

Our data show that *ubx5Δ* mutant cells are not sensitive to DPC induction by FA, HU (Figure 2A) nor CPT (Figure S2A). This suggests that, although Ubx5 is crucial for turnover of some DNA-bound proteins following exposure to genotoxins (Figures 5A and 5B), it might not be essential for removal of all kinds of adducts targeted by Wss1. Consistent with that proposal, some recent *in vitro* experiments using biochemical reconstitution of purified proteins revealed that SPRTN/Wss1 is able to cope with loosely folded DPCs without the need of p97/Cdc48 prior to proteolysis (Kroning et al. 2022). One could therefore propose that the reliance on Ubx5-Cdc48 unfolding activity depends on the characteristics of the DPC encountered. Thus, if Ubx5-Cdc48 becomes engaged on tightly folded DPCs, it might absolutely require proteolysis by Wss1 to finalize the extraction, which is demonstrated by a great Ubx5 accumulation at DPCs in the absence of Wss1 (Figure 3D). While Wss1 could be assisted by Ubx5-Cdc48 in a wide variety of DPCs, it may absolutely rely on Ubx5-Cdc48 unfolding activity only for a subset of tightly folded DPCs. This model could also partially explain why *wss1Δ* mutant cells complemented with a Wss1 allele deficient in Cdc48 binding are resistant to HU exposure (Maddi et al. 2020), but not to Top1-crosslink accumulation (Stingele et al. 2014).

### Cdc48 uses different adaptors to target different DPC types

The large variety of Cdc48 adaptors raises the question of how many can act in DPC repair pathways and whether they are redundant. Indeed, Cdc48/p97 can associate with its adaptors in different ways to form various assemblies (Alexandru et al. 2008; Schuberth and Buchberger 2008); Cdc48/p97 can also bind cofactors in a hierarchical manner to provide additional substrate specificity (Hanzelmann, Buchberger, and Schindelin 2011). Particularly, the mammalian UBXD7 protein (Ubx5 homolog) only binds to p97 in complex with UFD1-NPL4 and not to p97 alone (Hanzelmann, Buchberger, and Schindelin 2011). It is therefore not surprising to find both Ufd1 and Npl4 as suppressors of *tdp1Δ wss1Δ* mutant along with Ubx5 (Figure 1A). Similarly, SPRTN/Wss1 interacts specifically with p97/Cdc48 bound to UFD1-NPL4 (Davis et al. 2012). These data raise the possibility that Ubx5 acts in DPC repair specifically along with Cdc48-Ufd1-Npl4. The p97/Cdc48 adaptor TEX264 is strictly required for SPRTN-mediated repair of Top1ccs (Fielden et al. 2020) and is so far restricted to Top1cc repair only. Likewise, Ubx5 could be implicated in a specialized complex required for repair of a subtype of DPCs.

Among the UBX protein family, Ubx4 also turned out to be an interesting suppressor of *tdp1Δ wss1Δ* (Figures 1A and S2B). Given that Ubx4 and Ubx5 were redundant and in charge of a similar function in RNAPII turnover following UV damage (Verma et al. 2011), one could hypothesize a redundant role for Ubx4 and Ubx5 in DPC repair as well. This interpretation could explain why *ubx5Δ* shows no sensitivity towards DPC-inducing agents (Figures 2A and S2A), as Ubx4 could take over its role. Yet, our data do not exclude the potentiality of distinct substrate specificity for Ubx4 and Ubx5 in DPC repair.

To our surprise, the Cdc48 cofactor Doa1 did not appear as a suppressor of *tdp1Δ wss1Δ* in the transposon screen data from Serbyn et al. (2020) (Figure 1A). Indeed, Doa1 works in DPC repair along with Cdc48 and Wss1 in a ternary complex, capable of targeting SUMO substrates (Balakirev et al. 2015), although Doa1 was thought to be ubiquitin specific (Mullally, Chernova, and Wilkinson 2006). These conclusions were drawn from precipitation experiments performed without any stress conditions, and the function of Doa1 in DPC repair was not tested directly. Doa1 may also be involved in the repair of a specific subtype of DPC not tested in our study.

Altogether, these observations support the view that Cdc48/p97 assists Wss1/SPRTN in the repair of a wide range of DPCs, depending on the cofactor it is associated with. It remains to be understood how many Cdc48/p97 cofactors have a relevant function in DPC repair biology and in which specific context they might be required.

### An alternative role for Ubx5 in replisome disassembly

In eukaryotes, replisome disassembly relies on the recruitment of Cdc48 to ubiquitylated CMG by Ufd1-Npl4 (Maric et al. 2014; Maric et al. 2017). Recent observations in *Xenopus laevis* (*X. laevis*) suggested that Ubxn7 (Ubx5 ortholog) facilitates replisome disassembly by enhancing the recognition of ubiquitin-modified substrates by p97/Cdc48 (Tarcan et al. 2021). Similarly, UBXN-3, a member of the UBX family of CDC-48 cofactors in *Caenorhabditis elegans* (*C. elegans*), also stimulates CMG disassembly by CDC-48 (Franz et al. 2016; Sonneville et al. 2017; Xia et al. 2021). Although *in vitro* data showed that Cdc48-Ufd1-Npl4 is sufficient to disassemble ubiquitylated yeast CMG (Mukherjee and Labib 2019), the possibility that this critical process is more tightly regulated *in vivo* cannot be ruled out. Accordingly, one could hypothesize that yeast Ubx5 may control efficiency of replisome disassembly as well. In this view, the *ubx5Δ* suppression effect on *tdp1Δ wss1Δ* (Figure 1) may be attributed to reduced efficiency of replisome disassembly in these DPC-harboring mutant cells. In agreement with this possibility, previous work showed that DPCs can be repaired in a replication-dependent manner (Duxin et al. 2014; Vaz et al. 2016; Larsen et al. 2019). Particularly, the protease Ddi1 was proposed to reach a DPC-site in an S-phase dependent manner (Serbyn et al. 2020), consistent with Ddi1 travelling with the replication machinery. Since Ddi1 is crucial for repair of *ubx5Δ tdp1Δ wss1Δ* (Figures 4 and 5), we propose that the replisome may bring additional DPC repair factors such as Ddi1 to damage sites (Figure 4D). According to this model, it is likely that efficient replisome unloading, streamlined by Ubx5, would not leave enough time for efficient processing of the adduct in the absence of Wss1 and Tdp1. As a result, Ubx5 accumulation at a DPC locus (Figure 3D) would trigger too rapid and efficient replisome unloading, which may result in DPC proteolysis defects. Consistently, loss of Ubx5 would slow-down replisome disassembly, leaving more time for DPC proteolysis by alternative repair routes. We propose here an alternative role of Ubx5 in replisome unloading, emphasizing the critical importance of managing this process, which, if not tightly regulated, can lead to genome instability.

## MATERIALS AND METHODS

### Yeast strains and growth conditions

All *S. cerevisiae* yeast strains used in this study (listed in Table S1) were derived from W303 or S288C genetic backgrounds. Gene disruptions were made by homologous recombination using pFa6a PCR-based cassettes flanked by sequences adjacent to the gene to delete. Deletions were verified by colony PCR and phenotypic analyses. Epitope insertions were checked by immunoblotting (see *Protein extraction and Western blotting*). Other mutants were obtained by standard techniques and tetrad dissection (see *Genetic crosses and Tetrad dissection*).

Yeast cells were grown at 30 °C in YEP- (1 % yeast extract, 2 % peptone) or SC- (1.7 g/L yeast nitrogen base; 5 g/L ammonium sulfate; 0.87 g/L dropout mix) liquid media, or grown on plates supplemented with 20 g/L agar. 2 % glucose, 2 % raffinose or 2-3 % galactose was added as a source of sugar.

### Bacterial strains and growth conditions

DH5α *Escherichia coli* (*E. coli*) bacterial strains (listed in Table S2) were grown at 37°C in LB medium (1% bacto-tryptone, 0.5% yeast extract, 1% NaCl) or on LB-2% agar plates supplemented with 50 μg/mL of ampicillin for plasmid selection.

### Yeast transformation

5mL of exponentially growing cells were resuspended in 80 μL LiTE buffer (100 mM LiAc; 10 mM Tris pH 7.5; 1 mM EDTA). 38 μL of cells were mixed with 100 μg/mL salmon sperm ssDNA, 37.28 % PEG4000 and 200 ng DNA to transform (plasmid DNA or PCR product). Cells were incubated for 1 h at 30 °C, then supplemented with 6 % DMSO and a heat shock was performed for 10 min at 42 °C. Cells were plated on selective media and grown for 2-3 days before isolation of single colonies.

### Genetic crosses and Tetrad dissection

Haploid strains of opposite mating types were mixed and spotted overnight on a YEP-2% glucose (YEPD) plate at 30 °C, to allow mating and diploid construction. Diploids were selected by streaking the mating mixture overnight onto a plate selecting for the diploid genotype. Spore formation was induced by transferring the diploid cells to KAc sporulation medium (20 g/L potassium acetate; 2.2 g/L yeast extract; 0.5 g/L glucose; 0.87 g/L dropout mix; 20 g/L agar; pH 7) for 4-5 days at 30 °C. Before tetrad dissection, the ascus cell wall was digested with 0.5 mg/mL Zymolyase (Amsbio 120491-1) treatment for 5 min at room temperature. Haploid spores from single tetrads were separated with a micromanipulator and grown on a YEPD plate for 3 days at 30 °C. The individual phenotype of the dissected spores was determined by replica plating on agar plates containing selective media (various drop-out and drug media).

### Spot assays

Yeast mutant growth and sensitivities were assessed by spot assay. Cells were grown exponentially in the appropriate medium at 30 °C under continuous rotation and diluted to 0D_600_ = 1-1.5. 10-fold serial dilutions were spotted on agar plates containing the indicated concentrations of auxin, CPT or hydroxyurea HU. To treat cells with FA, 1 mL of yeast culture was incubated with FA for 15 min under rotation. Cells were then harvested by centrifugation, washed twice with 1 mL sterile water then diluted and spotted on YEPD plates.

### Protein extraction by TCA, Western Blotting and Antibodies

Standard trichloroacetic acid (TCA) extraction was used for analysis of total protein levels. Yeast cultures were fixed with 6.25 % of TCA, kept on ice for 10 min, pelleted by centrifugation at 5000 rpm for 10 min, washed twice with 1 mL of 100 % acetone and dried under vacuum. Pellets were resuspended in 100 μL of Urea Buffer (50 mM Tris-HCl pH 7.5; 5 mM EDTA; 6 M Urea; 1 % SDS), mixed with 200 μL of glass beads and subjected to bead-beating in a MagNa Lyzer instrument (Roche) 5 times for 45 secs at 4°C, 1 min of pause in between. Final solubilization was performed at 65°C for 10 min and centrifugation 10 min at 13000 rpm. Addition of 100 μL protein sample buffer (3 % SDS; 15 % Glycerol; 0.1 M Tris pH 6.8; 0.0133 % bromophenol blue; 0.95 M 2-mercaptoethanol) and boiled for 10 min. Protein samples were separated by SDS-PAGE and transferred onto nitrocellulose membrane. After transfer, membranes were blocked for 30 min with 5 % milk and incubated with appropriate antibodies indicated in each figure. See Table S3 for a list of antibodies used in this study.

### Flp-nick induction, Chromatin Immunoprecipitation (ChIP) and quantitative real-time PCR (qPCR)

Induction of the *flp-H305L-*3HA expression was performed as described in (Nielsen et al. 2009). Briefly, yeast cells were grown to log phase in YEP-2 % raffinose (no transcription of the locus). Where indicated, cells were synchronized in G1 with 200 ng/mL α-factor for 1.5 h. Induction was performed for 2 h by addition of 3 % galactose. Cells were then washed twice with 20 mL cold YEP (no sugar) medium and released into warm YEPD medium. 1 mL of cells were collected to monitor cell-cycle progression at desired timepoints (see *Cell cycle analysis by flow-cytometry analysis*).

Cells were harvested at the different timepoints by centrifugation at 3000 rpm for 3 min, washed twice with cold 1x Phosphate Buffer Saline (PBS) and frozen in 2 mL screw-cap tubes. For Ddi1-TAP, Ubx5-TAP, and Ubiquitin ChIP, cells were fixed before pelleting by addition of 1 % formaldehyde (FA) for 15 min at room temperature, then quenched with 250 mM glycine for 5 min and kept on ice for at least 10 min. All subsequent steps were performed at 4 °C. The frozen cells were resuspended in 1 mL of cold FA lysis buffer [50 mM HEPES-KOH, pH 7.5; 140 mM NaCl; 1 mM EDTA; 1 % Triton X-100; 0.1 % sodium deoxycholate; protease inhibitor cocktail (Roche)] and lysed by bead-beating with 500 μL of 0.5 mm glass beads with 5 cycles of 30 secs at 6000 rpm with 1 min pause in between each bead beating cycle, on a MagNA Lyser Instrument (Roche). Lysates were then recovered in a new tube by centrifugation and further spun at 13000 rpm for 30 min. Pellets were resuspended in 1 mL of FA lysis buffer and subjected to DNA sonication for 20 cycles of 30 secs in a Bioruptor Twin (Diagenode). Following pelleting by 15 min centrifugation at 13000 rpm, soluble fractions were transferred to a new tube and protein concentration was measured by Bio-Rad protein assay (500-0006). For each ChIP, 1 mg of protein (1/10 of input transferred in a new tube) was incubated together with 1 μL of anti-HA antibodies (BioLegend 901502 anti-HA.11 clone 16B12, for flp-H305L-3HA ChIP), or 1 μL of anti-Ubiquitin antibodies (Calbiochem ST1200-100UG clone FK2), or 20 μL of Dynabeads Pan Mouse IgG (Invitrogen 11041, for Ddi1-TAP and Ubx5-TAP ChIP) overnight at 4 °C with rotation. Prewashed Protein G Dynabeads (Invitrogen 100009D) were then added for 3 h to recover antibodies. Beads were collected on magnetic stands and washed once with 500 μL of FA lysis buffer, twice with 500 μL of FA-500 buffer (50 mM HEPES-KOH, pH 7.5 ; 500 mM NaCl ; 1 mM EDTA ; 1 % Triton X-100 ; 0.1 % sodium deoxycholate), twice with 500 μL of Buffer III (10 mM Tris-HCl pH 8 ; 1 mM EDTA ; 250 mM LiCl ; 1 % IGEPAL ; 1 % sodium deoxycholate) and finally once with 500 μL of TE buffer (50 mM Tris-HCl pH 7.5 ; 10 mM EDTA). Proteins were eluted twice with 100 μL of elution buffer B (50 mM Tris-HCl pH 7.5; 1 % SDS; 10 mM EDTA) for 8 min at 65 °C. Eluted and input samples were incubated for 2 h at 42 °C with 0.75 mg/mL Proteinase K (Carl Roth 7528.3) and decrosslinked at 65 °C for at least 12 h. DNA was purified with the MinElute PCR purification kit (Qiagen 28006) and eluted from the columns twice with 30 μL of elution buffer from the kit. Real-time qPCR reactions were then performed using SYBR Select Master Mix (Applied Biosystems 4472942) with oligonucleotide pairs listed in Table S4 (FRT +200 bp; +500 bp; +1000 bp or intergenic). Results are presented as percent of input or normalized to the intergenic region.

### Cell Cycle Analysis by Flow Cytometry Analysis (FACS)

For flow cytometry analysis, 1 mL of yeast cells at OD_600_ = 0.5 were harvested by centrifugation and resuspended in 70% ethanol allowing storage at 4°C for up to 1-2 weeks. Next, cells were pelleted at 6000 rpm for 2 min, washed with 300 μL of 50 mM Sodium Citrate (NaCi) pH 7.2. A second centrifugation at 6000 rpm for 10 min was performed, and cells were resuspended in 250 μL NaCi + 5 μL RNase A and incubated for 1 h at 37 °C. Staining with 25 μg/mL Propidium Iodide (PI) was performed at 37 °C for 1 h. Cells were then sonicated in a Bioruptor Twin (Diagenode) during 5 cycles of 5 secs, before flow cytometry analysis.

### Analysis of Rpb1 levels following Hydroxyurea Treatment

Overnight cultures were diluted to OD_600_ = 0.2. When cells reached OD_600_ = 0.8, HU (Bio Basic HB0528) was added to a final concentration of 0.2 M along with 100 μg/mL cycloheximide (CHX) (Sigma-Aldrich C7698) to prevent protein synthesis. Cultures collected at different time points were subjected to TCA protein extraction (see *Protein extraction by TCA)* and immunoblotted with anti-Rpb1 (BioLegend 664912 clone 8WG16) or anti-Pgk1 (Abcam ab113687 clone 22C5D8) antibodies. Fluorescent secondary antibodies (LI-COR 926-32210) were used for quantification analyses.

### Isolation of Chromatin

The protocol was adapted from (Kubota et al. 2012). 50 OD of yeast cells were harvested by spinning down cultures for 3 min at 3000 rpm and washed once with 1x cold PBS and frozen. Pelleted cells were resuspended in 1 mL of pre-spheroblasting buffer (100 mM PIPES/KOH, pH 9.4; 10 mM DTT; 0.1 % sodium azide) and incubated for 10 min at room temperature and centrifuged for 3 min at 4100 rpm. To induce spheroblasts formation, cells were resuspended in 1 mL spheroblasting buffer (50 mM KH_2_PO_4_/K_2_HPO_4_, pH 7.4; 0.6M Sorbitol; 0.1 mM DTT; 0.5 mg/mL Zymolyase, 2 % Glusulase) and incubated 30 min at 37 °C. Spheroblasts were harvested by centrifugation at 4100 rpm for 3 min and washed twice in 1 mL Wash buffer [20 mM KH_2_PO_4_/K_2_HPO_4_, pH 6.5; 0.6M Sorbitol; 1 mM MgCl_2_; 1 mM DTT; 20 mM β-glycerophosphate; 1 mM PMSF; protease inhibitors (Roche)]. Spheroblasts were resuspended in 200 μL of Wash Buffer and 1/10 of total cell extract was kept as input. Isolation of nuclei was obtained by overlaying spheroblasts on 1.4 mL 18 % Ficoll solution [18 % Ficoll; 20 mM KH_2_PO_4_/K_2_HPO_4_, pH 6.5; 1 mM MgCl_2_; 1 mM DTT; 20 mM β-glycerophosphate; 1 mM PMSF; 0.01% IGEPAL; protease inhibitor (Roche)] and tubes were centrifuged for 5 min at 6800 rpm. The supernatant was subjected to a second spin at 6800 rpm for 5 min, and a final spin at 12300 rpm for 20 min to obtain the nuclei in the pellet. Nuclei were resuspended in 200 μL of EB-X buffer (50 mM HEPES/KOH, pH 7.5; 100mM KCl; 2.5 mM MgCl2; 0.1 mM ZnSO4; 2 mM NaF; 0.5 mM Spermidine; 0.25% Triton X-100; 1 mM DTT; 20 mM β-glycerophosphate; 1mM PMSF; protease inhibitor cocktail) and lysed for at least 10 min on ice. The entire preparation was then transferred on 500 μL of EBX-S Buffer (EB-X buffer; 30 % sucrose) and spun at 12000 rpm for 10 min. Chromatin pellet was gently resuspended in 1 mL EB-X buffer and finally centrifuged at 9700 rpm for 2 min. The chromatin was resuspended in 30 μL of 1.5 X Bolt LDS Sample Buffer (Life Technologies 1074433) and analyzed by immunoblotting with antibodies indicated on each figure. The quality of the fractionation was controlled by immunoblotting against PGK1 (cytoplasm contamination; Abcam ab113687 clone 22C5D8) and H3 (chromatin extraction; Invitrogen Pa5-16183).

## DATA AVAILABILITY

Data are available from the corresponding authors upon request. Further information and requests for resources and reagents should be addressed to the lead contact.

## ACKNOWLEDGMENTS

We thank Helle Ulrich for sharing the parental auxin degron strain; Lotte Bjergbaek for the Flp-nick system and Takeo Usui for the *12geneΔ0HSR* strain. We are grateful for Geraldine Silvano’s technical assistance. We thank all the members of the Stutz laboratory for critical reading of the manuscript, comments, suggestions and discussions.

## AUTHOR CONTRIBUTIONS

Conceptualization: A.N., F.S. Methodology: A.N., N.S., I.B., F.S. Investigation: A.N., I.B., N.S. Data Curation: A.N. Writing – Original Draft: A.N. Writing – Review and Editing: A.N., N.S., F.S. Supervision: F.S. Funding acquisition: F.S.

## SUPPLEMENTAL INFORMATIONS

Supplemental Figures S1-S4.

Table S1 – Strains used in this study and comments on their construction.

Table S2 – Plasmids used in this study.

Table S3 – Antibodies used in this study.

Table S4 – Oligonucleotides used in this study.

---

**Supplementary Figure 1 (related to Figure 1).**
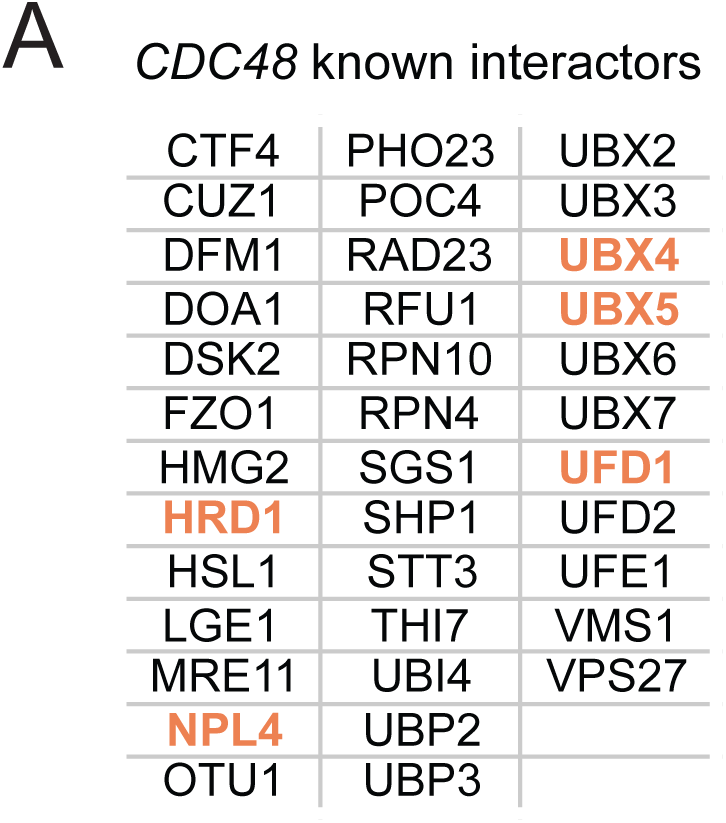
(A) List of known *CDC48* physical and genetic interactors extracted from SGD. The genes in orange correspond to *tdp1-degron wss1Δ* + auxin suppressors plotted in Figure 1A.

**Supplementary Figure 2 (related to Figure 2).**
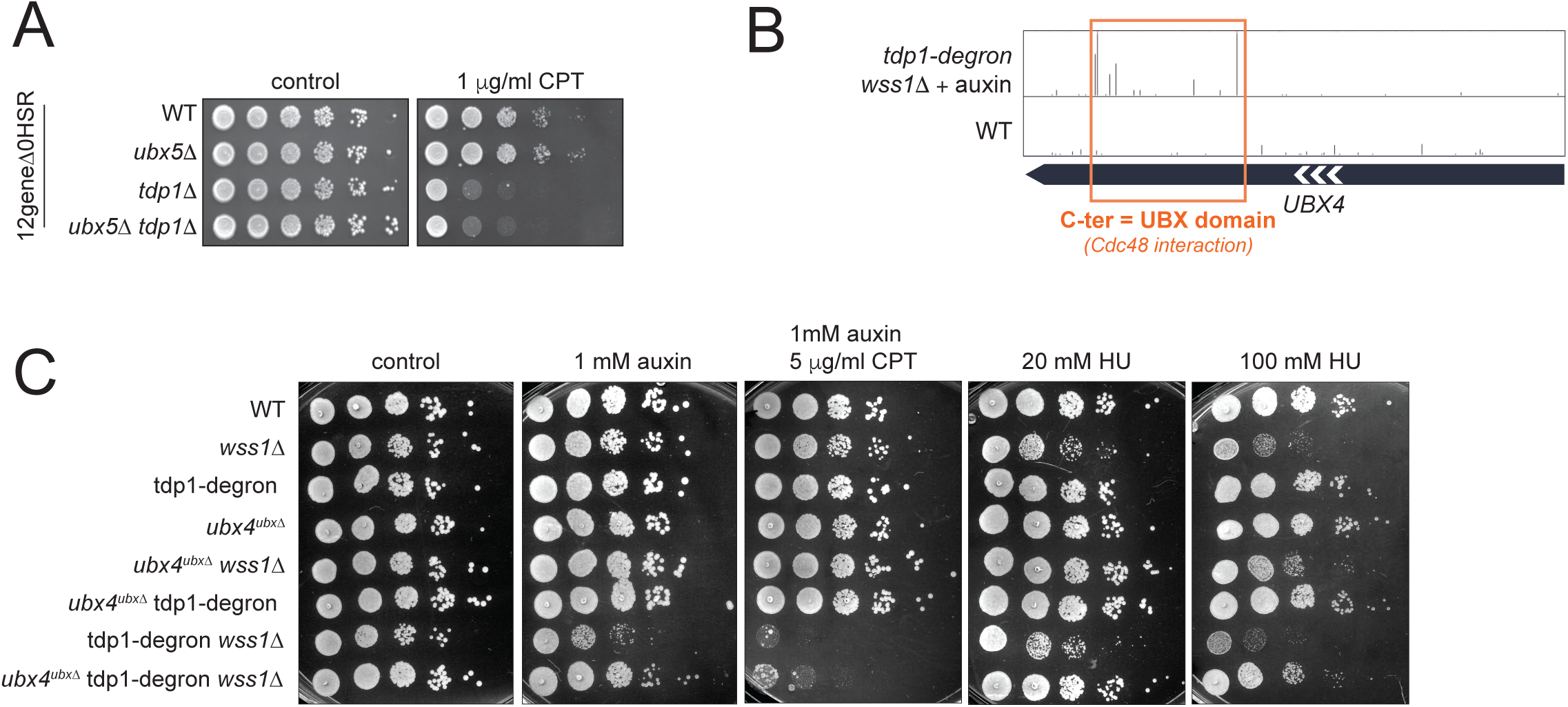
(A) *ubx5*1Δ is not suppressing defects linked to *tdp1*1Δ. The *12gene*1Δ*0HSR* mutant (Chinen et al. 2011) was used to reveal sensitivity to camptothecin (CPT). Cells were grown in YEPD and plated on 1 μg/mL CPT. Plates were incubated for 3 days at 30°C. (B) The C-ter UBX domain of Ubx4 is highly transposed in *tdp1wss1.* Snapshot depicting transposon coverage of *UBX4* gene body in the *tdp1-degron wss1Δ* + auxin and one of the WT libraries. The height of the bars represents the number of reads for each transposon. (C) Loss of the C-ter UBX domain of Ubx4 suppresses *wss1*1Δ and mildly suppresses *tdp1wss1*. Cells were grown in YEPD and spotted on a medium supplemented with 1 mM auxin, 5 μg/mL CPT, or indicated concentrations of HU. Plates were incubated for 2 days at 30°C.

**Supplementary Figure 3 (related to Figure 3).**
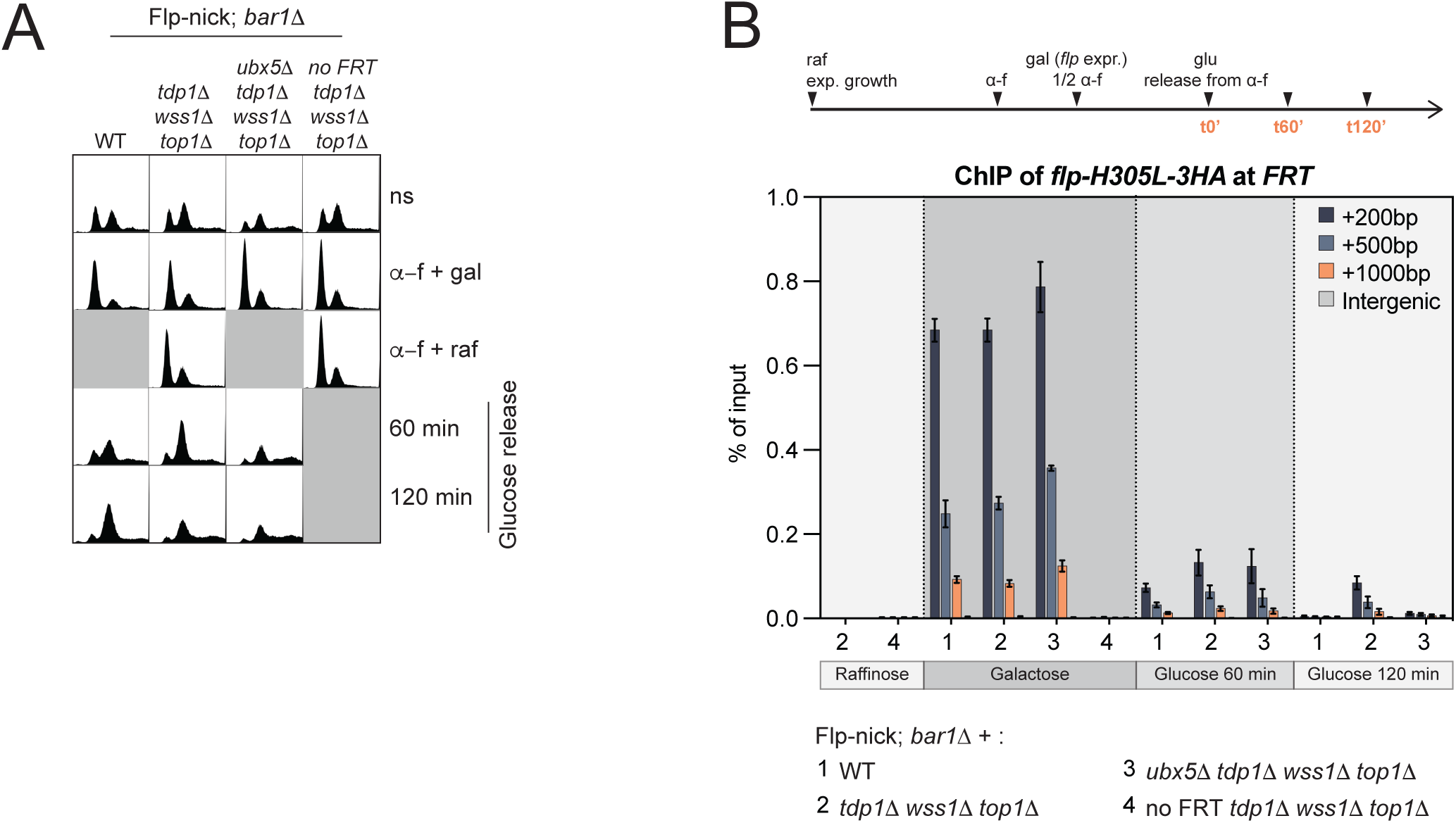
(A) Cell cycle progression of Flp-nick strains and time points used in Figure 3C, monitored by fluorescence-activated cell sorting (FACS). ns, non-synchronized; α-f, alpha-factor; gal, galactose; raf, raffinose. (B) *flp-H305L-3HA* levels at the *FRT* site presented as percent of input for the experiment described in Figure 3C.

**Supplementary Figure 4 (related to Figure 4).**
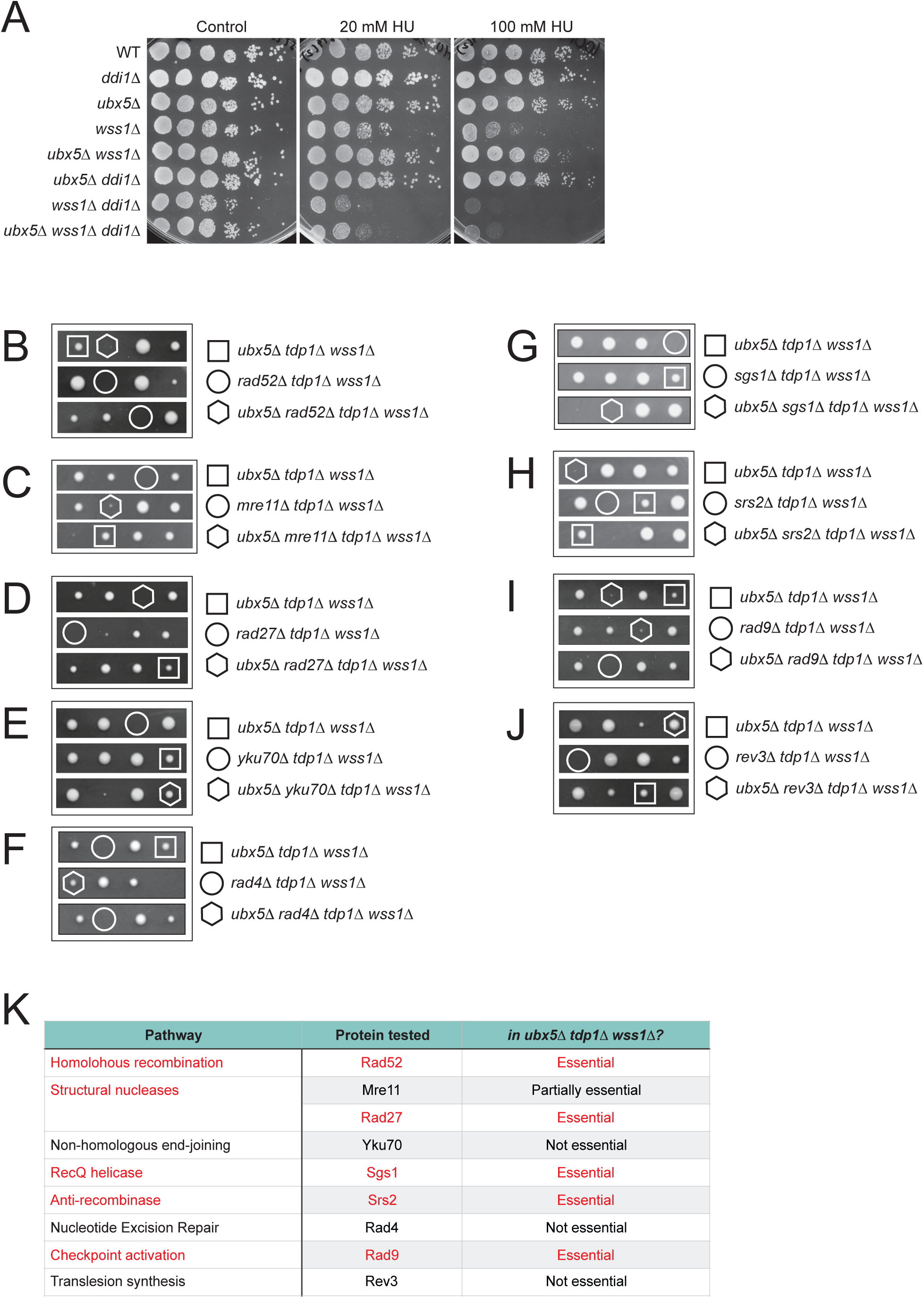
(A) *ubx5*1Δ restores *wss1*1Δ resistance to HU but not upon additional deletion of *DDI1.* Strains were grown in YEPD and spotted on the indicated concentrations of HU. Plates were incubated for 2 days at 30°C. (B-J) Genetic interactions of *rad52*1Δ (B), *mre111Δ* (C), *rad271Δ* (D), *yku701Δ* (E), *rad41Δ* (F), *sgs11Δ* (G), *srs21Δ* (H), *rad91Δ* (I) and *rev31Δ* (J) with *tdp11Δ, wss11Δ* and *ubx51Δ*. Tetrads were analyzed after dissection of the diploid [*TDP1/tdp1*1Δ; *WSS1/wss1*1Δ; *UBX5/ubx5*1Δ] in combination with one of the following: [*RAD52/rad52*1Δ]; [*MRE11/mre11*1Δ]; [*RAD27/rad27*1Δ]; [*YKU70/yku70*1Δ]; [*RAD4/rad4*1Δ]; [*SGS1/sgs1*1Δ]; [*SRS2/srs2*1Δ]; [*RAD9/rad9*1Δ]; [*REV3/rev3*1Δ]. (K) Summary table of all tetrad analyses. Red: negative genetic interactions.

