## Supplementary Table S1 for "Ubx5-Cdc48 assists the protease Wss1 at DNA-protein crosslink sites in yeast"

| | Internal Database Number | Genetic background | Genotype | Source | Generated by + comments |
|---|---|---|---|---|---|
| | FSY 4976 | W303 | W303-1A {MATa; his3-11_15; leu2-3_112; ura3-1; trp1D2; ade2-1; can1-100} | Euroscarf | N/A |
| | FSY 4977 | W303 | W303-1B {MATalpha; his3-11_15; leu2-3_112; ura3-1; trp1D2; ade2-1; can1-100} | Euroscarf | N/A |
| | FSY 7311 | BY4741 | MATa; his3Δ1; leu2Δ0; lys20; met15Δ0; ura3Δ0 | Euroscarf | N/A |
| 12geneD0HSR | FSY 8374 | BY4741 | MATa his3Δ1 leu2Δ0 met15Δ0 ura3Δ0 pdr3Δ0 pdr8Δ0 pdr1Δ0 yrr1Δ0 snq2Δ0 pdr5Δ0 pdr10Δ0 yor1Δ0 pdr15Δ0 pdr11Δ0 pdr12Δ0 aus1Δ0 RME1(ins308A) | Chinen et al., 2011 | N/A |
| Flp-nick Flp-HA | FSY7561 | W303 | Cir0; leu2-3_112::pGAL10-flp-H305L-3HA-HIS3; FRT-tetO-URA3; TetR-mRFP MATa | Serbyn et al., 2020 | N/A |
| Flp-nick Flp micr. | FSY 7400 | W303 | Cir0; leu2-3::pGAL-flp(H305L)-LEU2; TetR-mRFP(iYGL119W); tetOx224(iYFR020W)-URA3-FRT; Nup49-GFP | Serbyn et al., 2020 | N/A |
| **Figure 1 and Supplementary 1** | | | | | |
| **Figure 1C** | FSY 7625 - diploid | W303 | tdp1D::URA3/TDP1; wss1D::HIS3/WSS1; MATa/alpha | This study | cross - used for transformation and dissection (see below) |
| | FSY 7718 - diploid | W303 | tdp1D::URA3/TDP1; wss1D::HIS3/WSS1; ubx5D::KAN/UBX5; MATa/alpha | This study | transformation of PCR product ubx5D::KAN in the FSY 7625 diploid |
| **Figure 1D** | FSY 4976 | W303 | WT; MATa | Euroscarf | N/A |
| | FSY 7933 | W303 | ubx5D::KAN; MATa | This study | cross |
| | FSY 5859 | W303 | wss1D::HIS3; MATa | Serbyn et al., 2020 | N/A |
| | FSY 6229 | W303 | Tdp1-AID*-6HA-HPH; ura3-1::pADH-Os.TIR1-URA3. MATa | Serbyn et al., 2020 | N/A |
| | FSY 6263 | W303 | wss1D::HIS3; Tdp1-AID*-6HA-HPH; ura3-1::pADH-Os.TIR1-URA3; MATa | Serbyn et al., 2020 | N/A |
| | FSY 7935 | W303 | ubx5D::KAN; wss1D::HIS3; MATa | This study | cross |
| | FSY B1-17 | W303 | ubx5D::KAN; Tdp1-AID*-6HA-HPH; ura3-1::pADH-Os.TIR1-URA3; MATa | This study | cross |
| | FSY B1-19 | W303 | ubxD5::KAN; wss1D::HIS3; Tdp1-AID*-6HA-HPH; ura3-1::pADH-Os.TIR1-URA3; MATa | This study | cross |
| | FSY 7719 | W303 | | This study | used to obtain the strains above |
| **Figure 1E, 1F** | FSY 8481 + pFS 4218 | W303 | ubx5D::TRP1 (MATa) + pRS315-HA-UBX5 | This study | cross + plasmid transformation (see table S2) |
| | FSY 9321 + pFS 1336 | W303 | ubx5D::TRP1; wss1D::NAT; Tdp1-AID*-9MYC-HPH; ura3-1::pADH-Os.TIR1-URA3 (MATa) + pRS315 (empty) | This study | cross + plasmid transformation (see table S2) |
| | FSY 9321 + pFS 4218 | W303 | ubx5D::TRP1; wss1D::NAT; Tdp1-AID*-9MYC-HPH; ura3-1::pADH-Os.TIR1-URA3 (MATa) + pRS315-HA-UBX5 | This study | cross + plasmid transformation (see table S2) |
| **Figure S1B** | FSY 8232 | W303 | ubx4D::TRP1; MATalpha | This study | Transformation of FSY 4977 with ubx4D::TRP1 - used to cross with FSY 7835 |
| | FSY 7835 | W303 | ubx5D::KAN; wss1D::HIS3; tdp1D::URA3; MATa | This study | Used to cross with FSY 8232 |
| **Figure 2 and Supplementary 2** | | | | | |
| **Figure 2A** | FSY 4976 | W303 | WT; MATa | Euroscarf | N/A |
| | FSY 7933 | W303 | ubx5D::KAN; MATa | This study | cross |
| | FSY 5859 | W303 | wss1D::HIS3; MATa | Serbyn et al., 2020 | N/A |
| | FSY 7935 | W303 | ubx5D::KAN; wss1D::HIS3; MATa | This study | cross |
| **Figure 2C, 2D** | FSY 7935 + pFS 668 | W303 | ubx5D::KAN; wss1D::HIS3 (MATa) + pJG4-5-pGAL (empty) | This study | plasmid transformation (see table S2) |
| | FSY 7935 + pFS 4277 | W303 | ubx5D::KAN; wss1D::HIS3 (MATa) + pJG4-5-pGAL-HA-UBX5 | This study | plasmid transformation (see table S2) |
| | FSY 7935 + pFS 4278 | W303 | ubx5D::KAN; wss1D::HIS3 (MATa) + pJG4-5-pGAL-HA-uimD | This study | plasmid transformation (see table S2) |
| | FSY 7935 + pFS 4279 | W303 | ubx5D::KAN; wss1D::HIS3 (MATa) + pJG4-5-pGAL-HA-ubxD | This study | plasmid transformation (see table S2) |
| **Figure S2A** | FSY 8374 | 12geneD0HSR | see basic strains | Chinen et al., 2011 | N/A |
| | FSY 9296 | 12geneD0HSR | ubx5D::KAN; MATa | This study | transformation of ubx5D::KAN amplified by PCR in FSY 8374 |
| | FSY 8955 | 12geneD0HSR | tdp1D::HPH. MATa | Serbyn et al., 2021 | N/A |
| | FSY 9311 | 12geneD0HSR | ubx5D::KAN; tdp1D::HPH; MATa | This study | transformation of ubx5D::KAN amplified by PCR in FSY 8955 |
| **Figure S2B** | FSY 7626 | W303 | wss1D::HIS3/WSS1; Tdp1-AID*-6HA-HPH/TDP1; pADH-Os.TIR1-URA3/ura3-1; MATa/alpha | This study | Transformed with ubx4-ubxD-KAN and then dissected to obtain the spores above |
| **Figure 3 and Supplementary 3** | | | | | |
| **Figure 3B** | B1-43 | Flp-nick Flp-HA | bar1D (no marker); top1D::NAT; URA3-FRT; pGAL10-flp-H305L-3HA-HIS3; MATa | This study | cross |
| | B1-44 | Flp-nick Flp-HA | bar1D (no marker); wss1D::TRP1; tdp1D::HPH; top1D::NAT; URA3-FRT; pGAL10-flp-H305L-3HA-HIS3; MATa | This study | cross |
| | FSY 8781 | Flp-nick Flp-HA | bar1D (no marker); ubx5D::KAN; wss1D::TRP1; tdp1D::HPH; top1D::NAT; URA3-FRT; pGAL10-flp-H305L-3HA-HIS3; MATa | This study | cross |
| | FSY 8779 | Flp-nick Flp-HA no FRT | bar1D (no marker); ubx5D::KAN; wss1D::TRP1; tdp1D::HPH; top1D::NAT; pGAL10-flp-H305L-3HA-HIS3; MATa | This study | cross |
| | FSY 7597 x FSY 7599 | Flp-nick Flp-HA | N/A | This study | FSY 7597 and FSY 7599 were crossed and transformed with ubx5D::KAN ; positive transformants were selected by colony PCR and then diploids were dissected to select the strains abo |
| | FSY 7597 | Flp-nick Flp-HA no FRT | bar1D (no marker); wss1D::TRP1; tdp1D::HPH; top1D::NAT; pGAL10-flp-H305L-3HA-HIS3; MATalpha | This study | N/A |
| | FSY 7599 | Flp-nick Flp-HA | URA3-FRT (next to ARS607); bar1D (no marker); pGAL10-flp-H305L-3HA-HIS3; MATa | This study | URA3-FRT was amplified from a plasmid and transformed ; insertion was verified by colony PCR |
| **Figure 3C, S3A and S3B** | FSY 7599 | Flp-nick Flp-HA | bar1D (no marker); URA3-FRT; pGAL10-flp-H305L-3HA-HIS3; MATa | This study | cross |
| | B1-44 | Flp-nick Flp-HA | bar1D (no marker); wss1D::TRP1; tdp1D::HPH; top1D::NAT; URA3-FRT; pGAL10-flp-H305L-3HA-HIS3; MATa | This study | cross |
| | FSY 8781 | Flp-nick Flp-HA | bar1D (no marker); ubx5D::KAN; wss1D::TRP1; tdp1D::HPH; top1D::NAT; URA3-FRT; pGAL10-flp-H305L-3HA-HIS3; MATa | This study | cross |
| | FSY 8779 | Flp-nick Flp-HA no FRT | bar1D (no marker); ubx5D::KAN; wss1D::TRP1; tdp1D::HPH; top1D::NAT; pGAL10-flp-H305L-3HA-HIS3; MATa | This study | cross |
| **Figure 3D** | FSY 8925 | Flp-nick Flp micr. no FRT | Flp micr. No FRT (LEU2); bar1D::KAN; Ubx5-TAP-HIS3; tdp1D::HPH; wss1D::TRP1; top1D::NAT; MATa | This study | cross |
| | FSY 8921 | Flp-nick Flp micr. | Flp micr. (URA3/LEU2); bar1D::KAN; Ubx5-TAP-HIS3; MATa | This study | cross |
| | FSY 8923 | Flp-nick Flp micr. | Flp micr. (URA3/LEU2); bar1D::KAN; Ubx5-TAP-HIS3; tdp1D::HPH; wss1D::TRP1; top1D::NAT; MATa | This study | cross |
| | B1-53 | BY4741 | Ubx5-TAP-HIS3; MATa | Ghaemmarghami et al., 2003 | Used to amplify Ubx5-TAP by PCR |
| | FSY 7535 | Flp-nick Flp micr. | Flp micr. (URA3/LEU2); bar1D::KAN; top1D::NAT; wss1D::TRP1 | Serbyn et al., 2020 | used to cross with FSY 8826 and FSY 8827 to obtain the above mutants |
| | FSY 8826 | Flp-nick Flp micr. | Flp micr. (URA3/LEU2); wss1D::TRP1; tdp1D::HPH; top1D::NAT; Ubx5-TAP-HIS3; MATalpha | This study | obtained by transformation of Ubx5-TAP - used to cross with FSY 7535 |
| | FSY 8827 | Flp-nick Flp micr. no FRT | Flp micr. No FRT (LEU2); wss1D::TRP1; tdp1D::HPH; top1D::NAT; Ubx5-TAP-HIS3; MATalpha | This study | obtained by transformation of Ubx5-TAP - used to cross with FSY 7535 |
| **Figure 4 and Supplementary 4** | | | | | |
| **Figure 4A** | FSY 7836 | W303 | ubx5D::KAN; wss1D::HIS3; tdp1D::URA3; MATalpha | This study | used to cross with 7337 |
| | FSY 7337 | W303 | ddi1D::TRP1; MATa | Serbyn et al., 2020 | used to cross with 7836 |
| **Figure 4B, 4C** | FSY 7692 - diploid | W303 | Ddi1-13MYC-TRP1/DDI1; ubx5D::KAN/UBX5; Tdp1-AID*-6HA-HPH/TDP1; TIR1-URA3/ura3-1; wss1D::HIS3/WSS1; MATa/alpha | This study | Diploid issected to get the mutants for Ddi1-13MYC levels |
| **Figure 4D, 4E** | FSY 9337 | Flp-nick Flp-HA | bar1D (no marker); Ddi1-TAP-LEU2; wss1D::TRP1; tdp1D::HPH; top1D::NAT; URA3-FRT; pGAL10-flp-H305L-3HA-HIS3; MATa | This study | transformation of Ddi1-TAP-LEU2 |
| | FSY 9335 | Flp-nick Flp-HA | bar1D (no marker); ubx5D::KAN; Ddi1-TAP-LEU2; wss1D::TRP1; tdp1D::HPH; top1D::NAT; URA3-FRT; pGAL10-flp-H305L-3HA-HIS3; MATa | This study | transformation of Ddi1-TAP-LEU2 |
| | FSY B1-44 | Flp-nick Flp-HA | bar1D (no marker); wss1D::TRP1; tdp1D::HPH; top1D::NAT; URA3-FRT; pGAL10-flp-H305L-3HA-HIS3; MATa | This study | used for transformation |
| | FSY 8781 | Flp-nick Flp-HA | bar1D (no marker); ubx5D::KAN; wss1D::TRP1; tdp1D::HPH; top1D::NAT; URA3-FRT; pGAL10-flp-H305L-3HA-HIS3; MATa | This study | used for transformation |
| **Figure S4A** | FSY 4976 | W303 | WT; MATa | Euroscarf | N/A |
| | FSY 7337 | W303 | ddi1D::TRP1; MATa | Serbyn et al., 2020 | N/A |
| | FSY 7933 | W303 | ubx5D::KAN; MATa | This study | cross |
| | FSY 5859 | W303 | wss1D::HIS3; MATa | Serbyn et al., 2020 | N/A |
| | FSY 7935 | W303 | ubx5D::KAN; wss1D::HIS3; MATa | This study | cross |
| | FSY 7991 | W303 | ubx5D::KAN; ddi1D::TRP1; MATa | This study | cross |
| | FSY 8012 | W303 | wss1D::HIS3; ddi1D::TRP1; MATa | Serbyn et al., 2020 | N/A |
| | FSY 7992 | W303 | ubx5D::KAN; wss1D::HIS3; ddi1D::TRP1; MATa | This study | cross |
| | FSY 7835 | W303 | ubx5D::KAN; wss1D::HIS3; tdp1D::URA3; MATa | This study | used to cross with FSY 7337 to obtain the above mutants |
| **S4B - S4J** | FSY 7935 | W303 | ubx5D::KAN; wss1D::HIS3; tdp1D::URA3; MATa | This study | used for crosses |
| | FSY 7936 | W303 | ubx5D::KAN; wss1D::HIS3; tdp1D::URA3; MATalpha | This study | used for crosses |
| | FSY 9251 | W303 | rad52D::NAT; MATalpha | This study | crossed with FSY 7535 |
| | FSY 7124 | W303 | mre11D::HPH; MATa | Serbyn et al., 2021 | crossed with FSY 7536 |
| | FSY 5821 | W303 | rad27D::LEU2 MATalpha | Serbyn et al., 2021 | crossed with FSY 7535 |
| | FSY 9244 | W303 | yku70D::HPH MATa | This study | crossed with FSY 7536 |
| | FSY 7718 + rad4D::TRP1 - diploid | W303 | rad4D::TRP1/RAD4; tdp1D::URA3/TDP1; wss1D::HIS3/WSS1; ubx5D::KAN/UBX5; MATa/alpha | This study | FSY 7718 diploid was transformed with rad4D::TRP1 and dissected |
| | FSY 5863 | W303 | sgs1D::HPH; MATalpha | This study | crossed with FSY 7535 |
| | FSY 7381 | W303 | srs2D::TRP1; MATalpha | This study | crossed with FSY 7535 |
| | FSY 7373 | W303 | rad9D::HPH; MATalpha | This study | crossed with FSY 7535 |
| | FSY 9245 | W303 | rev3D::TRP1 MATalpha | This study | crossed with FSY 7535 |
| **Figure 5** | | | | | |
| **Figure 5A - 5D** | FSY 4976 | W303 | WT; MATa | Euroscarf | N/A |
| | FSY 7933 | W303 | ubx5D::KAN; MATa | This study | cross |
| | FSY 7935 | W303 | ubx5D::KAN; wss1D::HIS3; MATa | This study | cross |
| | FSY 7992 | W303 | ubx5D::KAN; wss1D::HIS3; ddi1D::TRP1; MATa | This study | cross |
| | FSY 5859 | W303 | wss1D::HIS3; MATa | Serbyn et al., 2020 | N/A |
| | FSY 7337 | W303 | ddi1D::TRP1; MATa | Serbyn et al., 2020 | N/A |
