## Supplementary Table S2 for "Ubx5-Cdc48 assists the protease Wss1 at DNA-protein crosslink sites in yeast"

### Plasmids used for transformation

| Internal Database number | | Source | Comments |
|---|---|---|---|
| pFS 668 | pJG4-5-pGAL (empty) | Gyuris et al., 1993 | N/A |
| pFS 4277 | pJG4-5-pGAL-HA-UBX5 | This work | Constructed by gibson assembly - sequence was checked by sequencing |
| pFS 4278 | pJG4-5-pGAL-HA-uimD | This work | Constructed by gibson assembly - sequence was checked by sequencing |
| pFS 4279 | pJG4-5-pGAL-HA-ubxD | This work | Constructed by gibson assembly - sequence was checked by sequencing |
| pFS 4218 | pRS315-HA-UBX5 | This work | Constructed by gibson assembly - sequence was checked by sequencing |
| pFS 1336 | pRS315 (empty) | Stutz lab stock | N/A |
