## Supplementary Table S3 for "Ubx5-Cdc48 assists the protease Wss1 at DNA-protein crosslink sites in yeast"

| Antibody | Company | Clone | Reference | RRID |
|---|---|---|---|---|
| anti-HA | Biolegend | 13B12 | 901502 | AB_2565006 |
| anti-Ubiquitin | calbiochem | FK2 | ST1200-100UG | AB_2043482 |
| anti-Rpb1 | Biolegend | 8WG16 | 664912 | AB_2650945 |
| anti-Pgk1 | Abcam | 22C5D8 | ab113687 | AB_10861977 |
| anti-MYC | Homemade | N/A | N/A | N/A |
| anti-Histone H3 | Invitrogen | | PA5-16183 | AB_10985434 |
| Fluorescent secondary Goat IRDye 800CW anti-Mouse | LI-COR | N/A | 926-32210 | AB_621842 |
| Secondary Goat anti-Mouse-HRP | DAKO | N/A | P0447 | AB_2617137 |
| Secondary Goat anti-Rabbit-HRP | DAKO | N/A | P0448 | AB_2617138 |
