## Supplementary Table S4 for "Ubx5-Cdc48 assists the protease Wss1 at DNA-protein crosslink sites in yeast"

| Name | Concentra | Source | Sequence |
|---|---|---|---|
| OFS4363_FRT +200bp_forward | 500 nM | Serbyn et al., 2020 | AAGTTCGACATGGGCTTCAG |
| OFS4364_FRT +200bp_reverse | 500 nM | Serbyn et al., 2020 | TCGTTTGGAGGACCTTTGAG |
| OFS4365_FRT +500bp_forward | 300 nM | Serbyn et al., 2020 | CGGGCAGTAGCTCATCAAGT |
| OFS4366_FRT +500bp_reverse | 300 nM | Serbyn et al., 2020 | CATGAAGAGGGTGAGGAGGA |
| OFS4367_FRT +1kb_forward | 600 nM | Serbyn et al., 2020 | CAGCCTGATCATTCATTCCA |
| OFS4368_FRT +1kb_reverse | 600 nM | Serbyn et al., 2020 | CGGACATCACAATCTTGCAC |
| OFS2788_intergenic_forward | 500 nM | Gali et al., 2017 | TGTTCCTTTAAGAGGTGATGGTGA |
| OFS2789_intergenic_reverse | 500 nM | Gali et al., 2017 | GTGCGCAGTACTTGTGAAAACC |
